# Structural basis of diverse antibody recognition of conserved coronavirus spike S2 epitopes that contribute to protective immunity

**DOI:** 10.64898/2026.09.13.750764

**Authors:** Adonis A. Rubio, Virginia Crivelli, Václav Hönig, Monika Čížková, Tomás Cervantes Rincón, Morgan E. Abernathy, Megan K. Parada, Danielle Vahdat, Yu E. Lee, Michael Eso, Jasmine Cantergiani, Simone Moro, David Jarrossay, Concetta Guerra, Benedetta Cena, Mihai Minculescu, Vaiva Gradauskaite, Maira Biggiogero, Veronica Calvaruso, Alessandra F. Pellanda, Andrea Cavalli, Christian Garzoni, Martin Palus, Davide F. Robbiani, Christopher O. Barnes

## Abstract

Conserved epitopes within the coronavirus spike S2 domain elicit broadly reactive antibodies, yet many characterized responses show limited neutralizing and variable protective activity, leaving their contribution to antiviral immunity unclear. Building on our previous mapping of evolutionarily conserved spike “coldspots”, we isolated human monoclonal antibodies targeting four conserved epitopes in the spike S2 domain: the internal fusion peptide (iFP), the central helix (CH), the connector domain (CD), and a membrane-proximal epitope in the heptad repeat 2 that we term the lower stalk (LS). A crystal structure of an LS-directed antibody defined a previously unresolved mode of antibody recognition of this membrane-proximal epitope, while cryogenic electron microscopy (cryo-EM) structures revealed that genetically diverse CH-specific antibodies use distinct binding modes to converge on conserved features of the prefusion S2 apex. Despite minimal neutralizing activity, CH- and LS-directed antibodies exhibited distinct antiviral functions. LS-directed antibodies mediated Fcγ receptor-dependent effector activity *in vitro*, whereas the broadly reactive CH-directed antibody ch.007 lacked detectable antibody-dependent cellular cytotoxicity (ADCC) or cellular phagocytosis (ADCP) activity yet protected mice from lethal SARS-CoV-2 MA10 challenge, with protection abrogated by Fcγ receptor-silencing mutations. Together, these findings expand the genetic, structural and functional landscape of human antibody responses to conserved coronavirus S2 epitopes and demonstrate that CH-directed antibodies can contribute to protective immunity through Fc-dependent mechanisms not predicted by *in vitro* neutralization or conventional *in vitro* Fc effector assays.

## INTRODUCTION

Coronaviruses (CoVs) are a family of enveloped, positive-sense single-stranded RNA viruses that infect a range of hosts including rodents, birds, pigs, camels, bats, and humans (*1, 2*). Their capacity for mutations and recombination events (*2*), positions CoVs as a persistent source of zoonotic spillover, underscoring the need for continued surveillance and preparedness. Currently, there are seven CoVs that are known to infect the human population: hCoV-229E, -NL63, -OC43, -HKU1, SARS-CoV, MERS-CoV, and SARS-CoV-2 (*3*). While four of these (hCoV-229E, -NL63, -OC43, and -HKU1) cause mild, seasonal respiratory illness, the impact of the SARS-CoV-2 pandemic highlights the threat that zoonotic CoV spillovers can have on the global economy and healthcare systems. Therefore, there is a need to identify and develop prophylactics and treatments that provide broad protection against the divergent CoV strains.

CoV-specific antibodies primarily target the spike glycoprotein, a trimeric class I fusogen on the viral surface responsible for attachment to the host cell receptors and subsequent membrane fusion (*4–8*). Upon cleavage by host proteases, spike is divided into the S1 and S2 subunits. S1 is comprised of the N-terminal domain (NTD) and receptor-binding domain (RBD) while the S2 subunit maintains the fusion peptide (FP), heptad repeat 1 (HR1), central helix (CH), connector domain (CD), and heptad repeat 2 (HR2) (*4–9*). The S1 subunit is immunodominant, with antibodies elicited during CoV exposure primarily targeting this region over the S2 (*10–12*). Throughout the SARS-CoV-2 pandemic, exclusively S1-specific monoclonal antibodies were approved for treatment given their potency at blocking virus entry and efficacy (*13, 14*). However, while immunodominant, the S1 subunit sequence is not highly conserved across SARS-CoV-2 variants and CoV strains (*15–17*), limiting the clinical deployment of S1-targeting antibodies. In contrast, the greater conservation of the S2 domain is attractive for its potential to elicit cross-reactive responses, as observed for other class I viral fusogens, such as those of influenza (*18–20*) and HIV-1 (*21, 22*).

The functional consequences of S2-directed antibodies are generally less well understood. Compared with antibodies targeting the S1 subunit, S2-directed monoclonal antibodies are generally less potent virus neutralizers but often exhibit greater cross-reactivity to diverse CoVs due to the higher sequence conservation of the fusion machinery. Broadly reactive antibodies have now been described against several conserved S2 regions, most prominently the FP (*23–29*), HR2 stem helix (SH) (*23, 30–36*), CH (*37–40*), and less prominently, the HR2 stalk (*41–43*). Whereas FP- and SH-specific antibodies can neutralize diverse coronaviruses and protect in animal models (*23–26, 32–36*), CH-directed and HR2-stalk antibodies characterized to date are generally weakly or non-neutralizing, and have not shown protection *in vivo*, although some CH-directed antibodies can engage Fc-dependent antiviral functions *in vitro* (*37–40*). Recent studies further suggest that portions of the S2 apex can elicit highly recurrent public antibody responses following SARS-CoV-2 exposure (*40*). Together, these findings suggest substantial functional heterogeneity among antibodies recognizing the conserved S2 core, but the genetic and structural diversity of these responses that can contribute meaningfully to antiviral protection through both virus neutralization-dependent and -independent mechanisms, remain unclear.

We previously used evolutionary sequence conservation to identify highly conserved, antibody-accessible “coldspots” within spike and demonstrated that conserved epitopes in the FP and SH are targeted by broadly neutralizing human antibodies with protective activity *in vivo* (*23*). However, whether additional S2 coldspots are accessible to human antibodies, how these responses recognize native spike, and whether antibodies with limited neutralizing activity can nevertheless contribute to protection remained unresolved. Here, we extend this framework by defining human monoclonal antibodies targeting four coldspots in the CoV S2: the internal fusion peptide (iFP) (*8, 44, 45*), the central helix (CH) (*4, 8, 9*), the connector domain (CD) (*4, 7, 8, 46, 47*), and the lower stalk (LS), a helical segment within HR2. Functional and structural characterization revealed that genetically diverse CH- and LS-specific antibodies display broad cross-reactivity across divergent CoVs, including zoonotic strains, and engage distinct antiviral functions despite minimal neutralizing activity *in vitro*. High-resolution structures further revealed multiple solutions for antibody recognition of the conserved CH, while *in vivo* studies demonstrated Fcγ receptor-dependent protection by the CH-directed antibody ch.007 despite the absence of detectable ADCC or ADCP activity *in vitro*. Collectively, these findings expand the genetic, structural and functional understanding of human antibody recognition across the conserved S2 subunit and highlight the diverse mechanisms through which S2-directed antibodies can contribute to antiviral immunity.

## RESULTS

### Human antibodies recognize evolutionarily stable coldspot epitopes in the CoV S2

To assess the stability of previously identified coldspots (*23*) in the context of ongoing SARS-CoV-2 viral evolution, we analyzed spike sequences after including those that became available at GISAID between April 2022 and July 2025 (n=17,417,909 total, see Methods). While some of the coldspots in S1 were truncated or shifted due to accumulated mutations, all previously defined S2 coldspots were confirmed (with the exception of a two amino acid (aa) shortening at the SH coldspot), demonstrating the sustained evolutionary stability of S2 coldpots despite extensive virus evolution (Figure S1A and Table S1). Furthermore, we added a new S2 coldspot of only 14 aa (cs16, residues 1177-1190), located in the HR2 domain, which we refer to as the lower stalk (LS) (Figure S1A and Table S1).

To determine whether these conserved epitopes elicit antibody responses during SARS-CoV-2 infection, we evaluated plasma from COVID-19 convalescent individuals (n = 71; Lugano cohort (*48*)) for IgG binding to peptides corresponding to each coldspot by the enzyme-linked immunosorbent assay (ELISA) (Figure 1A-C and S1B, Table S1). While little-to-no reactivity was observed against S1 coldspots, plasma IgG responses against multiple S2 coldspots were detected in at least a subset of donors (Figure 1A-C and S1B), including to the previously characterized FP and HR2 stem helix (SH) coldspots (*23*). Among responders, individuals with the highest plasma IgG reactivity to the remaining S2 coldspots were selected for monoclonal antibody isolation from peripheral blood B cells (Figure 1C-1D and S1C). Some of the antibodies were clonally related (Data File S1). In total, we recombinantly expressed 22 monoclonal antibodies: 2 targeting cs12 and 5 targeting cs13 (corresponding to the internal Fusion Peptide (iFP)), 7 targeting cs14 (central helix (CH)), 1 targeting cs15 (connector domain (CD)), and 7 targeting cs16 (lower stalk (LS)) (Data File S2).

**Figure 1.**
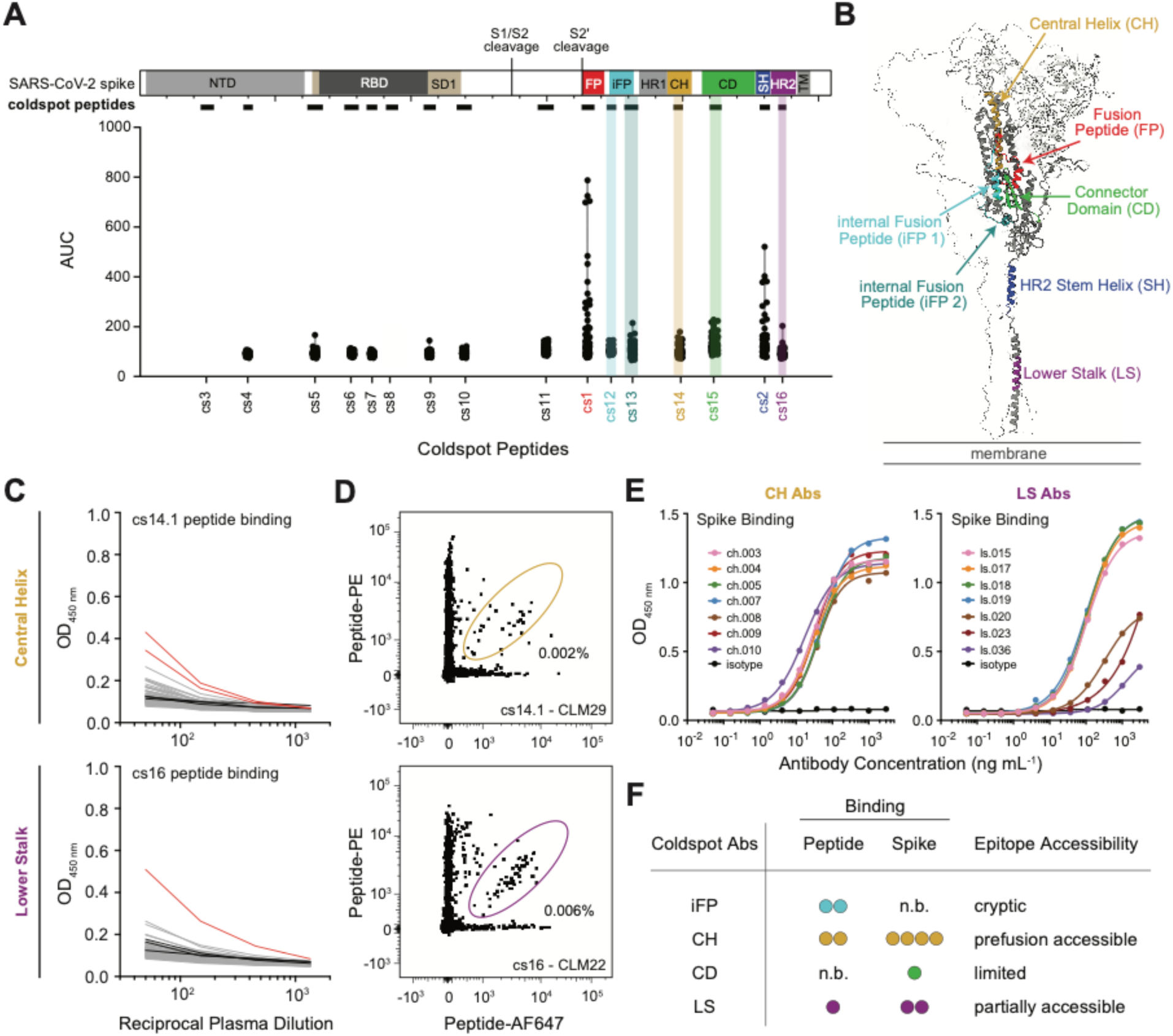
Identification of human antibodies targeting conserved CoV spike S2 coldspots. **(A)** Summary of plasma IgG reactivity to peptides corresponding to spike coldspots. Top: schematic of the SARS-CoV-2 spike with annotated domains and cleavage sites. Coldspot peptides are denoted by horizontal rectangles below the schematic (see Table S1). Bottom: cohort plasma IgG reactivity to each coldspot peptide represented as area under the curve (AUC) of the ELISAs in panel (C) and Figure S1B. **(B)** AlphaFold 3 (*55, 56*) model of the SARS-CoV-2 spike ectodomain (UniProt P0DTC2, residues 14-1208). One protomer is represented as cartoon, while the other two as surface. S2 coldspots from panel (A) are denoted in their respective colors. **(C)** Representative ELISA plots for plasma IgG reactivity to central helix (top) and lower stalk (bottom) coldspots in the S2 domain. Each grey or red line represents a convalescent individual (see Methods); red lines indicate individuals whose peripheral blood mononuclear cells (PBMCs) were selected for antigen-specific B cell sorting; black lines indicate pre-pandemic controls. See Figure S1B for reactivity to all other coldspots. Note: a peptide corresponding to cs3 could not be synthesized. **(D)** Representative flow cytometry plots of B cells binding to fluorescently labeled central helix (top) and lower stalk (bottom) peptides. Gating of double-positive cells, with respective percentages, are indicated within each plot. Peptide and donor sample labeled on the bottom right of each plot. See Figure S1C for full gating strategy. **(E)** Representative ELISA curves depicting central helix (left) and lower stalk (right) monoclonal antibodies’ binding to SARS-CoV-2 spike 2P trimer (see Figure S2A for binding of internal fusion peptide and connector domain antibodies and S2B for corresponding CH and LS peptide binding). **(F)** Summary matrix of monoclonal antibodies binding properties and epitope accessibility for internal fusion peptide (iFP), central helix (CH), connector domain (CD), and lower stalk (LS) coldspots.

By ELISA, the isolated monoclonal antibodies bound their cognate coldspot peptide and/or the full-length SARS-CoV-2 spike protein (2P stabilized; Figure 1E and S2A-B). While two iFP-specific antibodies bound their cognate peptide, neither recognized the recombinant spike protein (Figure S2A), suggesting limited accessibility of the iFP epitope within the prefusion trimer. Similarly, the CD-specific antibody exhibited weak binding to its cognate peptide and limited reactivity with spike (Figure S2A). Although iFP- and CD-directed antibodies did not exhibit detectable binding to soluble spike by ELISA, low but reproducible binding above the isotype control was observed by flow cytometry using cell-surface expressed spike (fig. S2D). In contrast, CH-specific antibodies bound their cognate peptide weakly (Figure S2B), yet recognized spike robustly, with EC_50_ values ranging from 7 to 90 ng mL^−1^ (Figure 1E and Data File S2). These findings are consistent with CH-directed antibodies recognizing structural or conformational features within the spike trimer that are not fully recapitulated by their linear peptide. LS-directed antibodies also demonstrated relatively weak binding to their cognate peptide (Figure S2B), but four antibodies–ls.015, ls.017, ls.018, and ls.019–bound spike with comparable EC_50_ values (104, 110, 107, and 103 ng mL^−1^, respectively) (Figure 1E and Data File S2).

We next assessed whether spike engagement with recombinant angiotensin-converting enzyme 2 (ACE2) influenced antibody binding, as previously reported for other S2-directed antibodies (*23, 25*). Using flow cytometry, we assessed antibody binding to native SARS-CoV-2 spike expressed on the surface of HEK293 cells, in the presence or absence of soluble ACE2 (Figure S2C-D). While some antibodies exhibited modest increases in the geometric mean fluorescence intensity (gMFI) upon engagement of ACE2, none showed the marked enhancement observed for fp.006, a FP antibody (*23*) (Figure S2C-D). Finally, evaluation in a SARS-CoV-2 pseudovirus neutralization assay (*49*), demonstrated that while several antibodies exhibited weak neutralizing activity at high concentrations, none were capable of potent pseudovirus neutralization (Figure S2E).

Together, these findings expand the set of human monoclonal antibodies targeting conserved S2 coldspots, including the trimeric spike accessible CH and LS epitopes (Figure 1F). While these antibodies exhibited minimal neutralizing activity, the accessibility and evolutionary conservation of these epitopes, combined with prior evidence that non-neutralizing antibodies can mediate protection against diverse viral pathogens (*50–54*), motivated further characterization of these antibody classes.

### Central Helix and Lower Stalk antibodies are cross-reactive with zoonotic CoVs

Sequence analysis revealed that the CH epitope is highly conserved across diverse betacoronaviruses, whereas the LS epitope exhibits somewhat lower but still substantial conservation, particularly among sarbecoviruses (Data File S3). Consistent with this conservation, CH-directed antibodies displayed broad binding reactivity across a diverse panel of human and zoonotic CoV spike proteins by ELISA (Figure 2A). Relative to the previously described CH antibody 54043-5 (*37*), ch.007 and ch.010 exhibited the greatest binding breadth, recognizing spike proteins from multiple divergent betacoronaviruses (Figure 2A). LS-directed antibodies also displayed cross-reactive binding, particularly within the sarbecovirus subgenus, with ls.015 and ls.019 exhibiting the strongest overall reactivity (Figure 2A, full data not shown).

**Figure 2.**
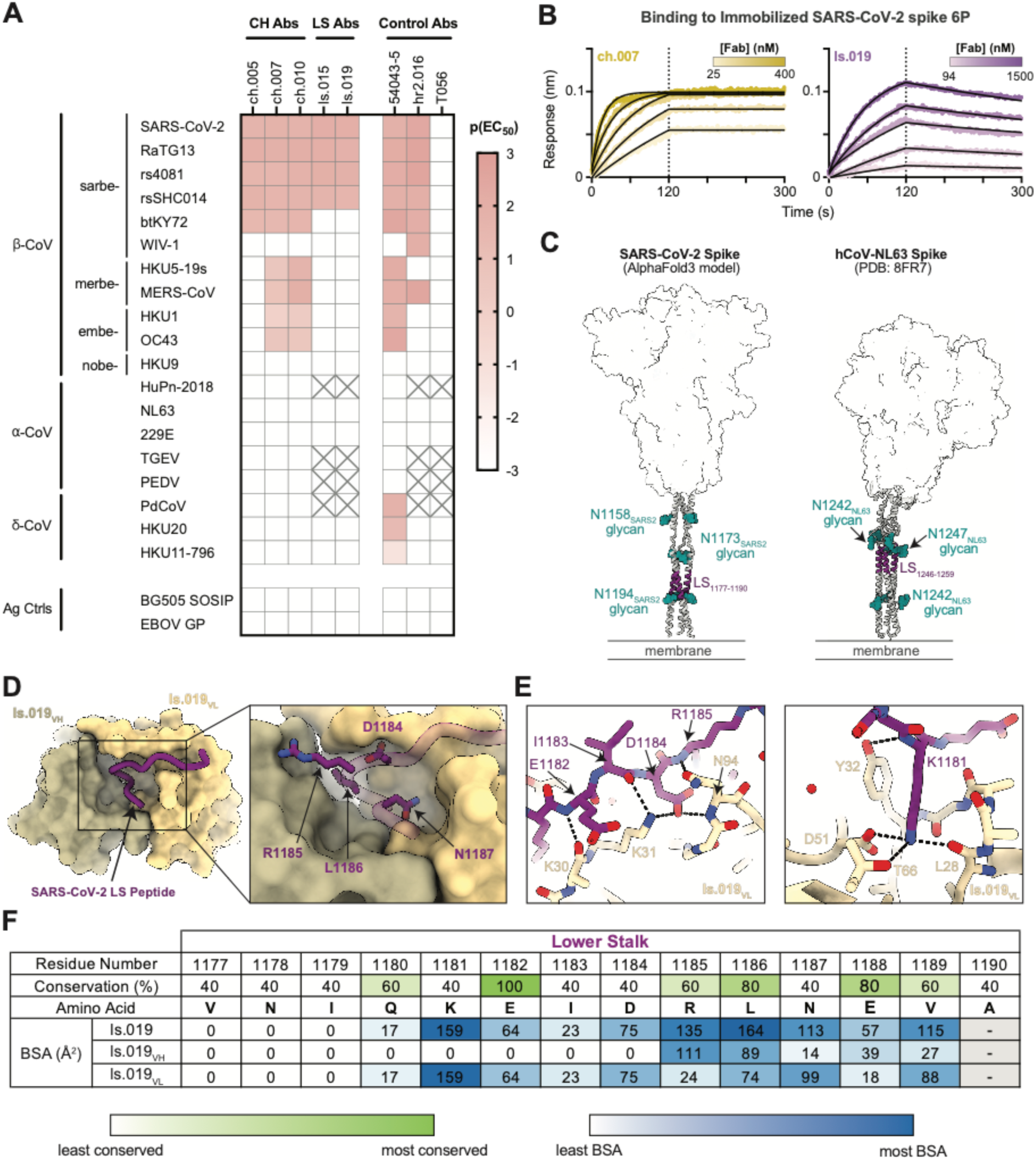
Cross-reactive antibodies target the conserved central helix and lower stalk S2 epitopes. **(A)** Heatmap depicting binding breadth of select CH (left) and LS (middle) antibody IgGs against SARS-CoV-2 spike 6P and related CoV spike 2P proteins, relative to negative control antigens (bottom) and antibodies (right). Controls include BG505 MD39 (*69*) v3.2 SOSIP, Ebola Zaire glycoprotein (GP), and 54043-5 (*37*), hr2.016 (*23*) and T056 (*70*) IgGs. **(B)** Representative sensorgrams from biolayer interferometry experiments depicting binding kinetics of ch.007 and ls.019 Fabs to immobilized SARS-CoV-2 spike 6P. Dotted line indicates start of dissociation step. **(C)** Depiction of the predicted helical conformation of the SARS-CoV-2 LS in the prefusion state, modeled using AlphaFold 3 (*55, 56*), compared to the experimentally modeled LS in the cryogenic electron tomography (cryo-ET) structure of hCoV-NL63. N-linked glycans are depicted by green spherical atoms; glycans for SARS-CoV-2 were modeled onto the AlphaFold structure using GlycoShape (*71*). **(D-E)** Visualization of molecular interactions between ls.019 scFv and the SARS-CoV-2 LS peptide, as elucidated by a 1.8 Å crystal structure. **(D)** The LS peptide is cradled by the ls.019 light chain and buries into a pocket formed at the interface of the ls.019 heavy and light chains. **(E)** Key hydrogen bonding between the peptide and Fab are represented by dashed lines. **(F)** Per residue contributions of the ls.019 V_H_ and V_L_ to the paratope buried surface area (BSA). Conservation of the LS peptide was calculated from a sequence alignment of representative betacoronaviruses: hCoV-OC43, hCoV-HKU1, SARS-CoV, SARS-CoV-2, and MERS-CoV.

To further define the molecular basis of this breadth, we examined the genetic features of the isolated antibodies. Whereas recently described antibody responses to the S2 apex can exhibit germline-restricted public responses that dominate recognition of portions of the S2 apex, the CH-directed antibodies here use diverse heavy and light chain V genes and lacked a shared heavy chain complementarity determining region 3 (CDRH3) signature (Data File S2), expanding the known genetic diversity of this antibody class and suggesting previously unrecognized modes of epitope engagement. Analysis of CDR3 lengths revealed notable deviations from the naïve B cell receptor (BCR) repertoire which is on average 13-15 aa at the heavy chain CDR3 (CDRH3) (*57–59*) (although SARS-CoV-2 specific BCR lengths are slightly skewed to ∼15-20 aa (*60–62*)) and ∼8-12 aa at the light chain CDR3 (CDRL3) (*58*). The two most cross-reactive CH-specific antibodies, ch.007 and ch.010, possess unusually short CDRH3, with lengths of 9 and 8 aa, respectively (Data File S2). In contrast, all LS-directed antibodies shared identical V-gene pairings (*IGHV3-30*03/IGLV3-27*01*) and CDR3 lengths, suggesting derivation from a common clonal lineage (Data Files S1-S2). These contrasting genetic profiles (i.e. diversity among CH-directed antibodies and clonal homogeneity among LS-directed antibodies) suggest distinct repertoire features associated with antibody responses to these conserved S2 epitopes.

To further quantify binding interactions with prefusion spike, we measured Fab binding kinetics to SARS-CoV-2 spike 6P (*63*) using biolayer interferometry (BLI) (Figure 2B). The broadly cross-reactive CH antibody ch.007 bound full-length spike with strong affinity despite targeting an epitope that is partially occluded by the S1 subunit in the prefusion trimer (Figure 2B). These findings are consistent with previous observations of transient “breathing” within the spike trimer (*46, 64, 65*). The ls.019 antibody also bound SARS-CoV-2 spike 6P appreciably, with a K_d_ value of 114 nM, albeit with a comparatively rapid dissociation rate (Figure 2B and Table S2).

Together, these data reveal that antibodies targeting the conserved CH and LS epitopes can recognize spike proteins from divergent coronaviruses. Whereas CH-directed antibodies arise from diverse genetic backgrounds and exhibit strong breadth across zoonotic betacoronaviruses, LS-directed antibodies represent a previously under-characterized antibody class with preferential reactivity toward sarbecoviruses.

### Structural characterization of the HR2 Lower Stalk epitope

To define the molecular basis of antibody recognition of the LS epitope, we determined a 1.8 Å crystal structure of the ls.019 single chain variable fragment (scFv) complexed with the LS peptide (SARS-CoV-2 residues 1177-1190) (Figure 2C-F and Table S3). To our knowledge, only one prefusion coronavirus structure has resolved the complete HR2 domain in the prefusion state: NL63 (PDB 8FR7) (*66*). Based on this structure and the AlphaFold3 model of the SARS-CoV-2 spike protein (Figure 2C), we anticipated the LS epitope to adopt a helical conformation. Unexpectedly, rather than adopting a helical conformation, the antibody-bound LS peptide adopted an extended conformation and was cradled within the ls.019 combining site (Figure 2D). Within this complex, the ls.019 V_L_ primarily engaged the N-terminal residues of the peptide, while the C-terminal segment, namely D1184_SARS-CoV-2_, L1185_SARS-CoV-2_, L1186_SARS-CoV-2_, and N1187_SARS-CoV-2_, was buried within a pocket formed by both the ls.019 V_H_ and V_L_ domains (Figure 2D-E). Further examination of the molecular interactions between residues also revealed key hydrogen bonding mediated between K1181_SARS-CoV-2_ and the light chain: L28_LC_ and Y32_LC_ in the CDRL1, D51_LC_ in the CDRL2, and T66_LC_ in the framework region 3 (Figure 2E). Analysis of both the sequence conservation and buried surface area (BSA) confirmed that while contacts between ls.019 and both K1181_SARS-CoV-2_ and L1186_SARS-CoV-2_ make up the majority of the paratope, K1181_SARS-CoV-2_ is not highly conserved across betacoronaviruses (Figure 2F and Data File S3). However, the LS is highly conserved among sarbecoviruses, with the bat CoV btKY72 being the most divergent (Data File S3), likely explaining the limited binding breadth of the LS antibodies (Figure 2A).

Although antibodies (*41–43*) and nanobodies (*67, 68*) targeting overlapping HR2 residues within the LS epitope have been described, our structure provides, to our knowledge, the first high-resolution molecular view of a human-derived antibody recognizing this conserved membrane-proximal HR2 epitope.

### Central Helix antibodies use distinct structural solutions to recognize conserved epitopes in the S2 apex

To define the structural basis of CH antibody recognition, we performed single particle cryo-EM of ch.005, ch.007, ch.008, and ch.010 Fabs bound individually to a prefusion-stabilized SARS-CoV-2 S2 stem (SS) construct incorporating stabilizing mutations from the HexaPro-S2 and HexaPro-SS designs (*72*) (Figure 3, S3-S4 and Table S4). We obtained global reconstructions of the four Fab-SS complexes at 3.2 Å, 2.6 Å, 3.2 Å and 3.0 Å resolution, respectively (Figure 3A-C, S3, and S4A-C). Consistent with previous observations of S2 “breathing”, in which the trimer apex transiently opens (*72–74*), all four maps displayed antibodies bound to a slightly open S2 conformation (Figure 3A-C, S3, and S4A-C). To improve local map features, we conducted focused refinements around each Fab-SS protomer interface (Figure 3A-C, S3, and S4A-C). Structural models were built for ch.005-, ch.007-, and ch.010-SS complexes, but not for ch.008-SS due to lack of confidence in assigning backbone density. Additionally, an X-ray structure for ch.005 Fab aided model building in the ch.005-SS protomer cryo-EM map (Table S3).

**Figure 3.**
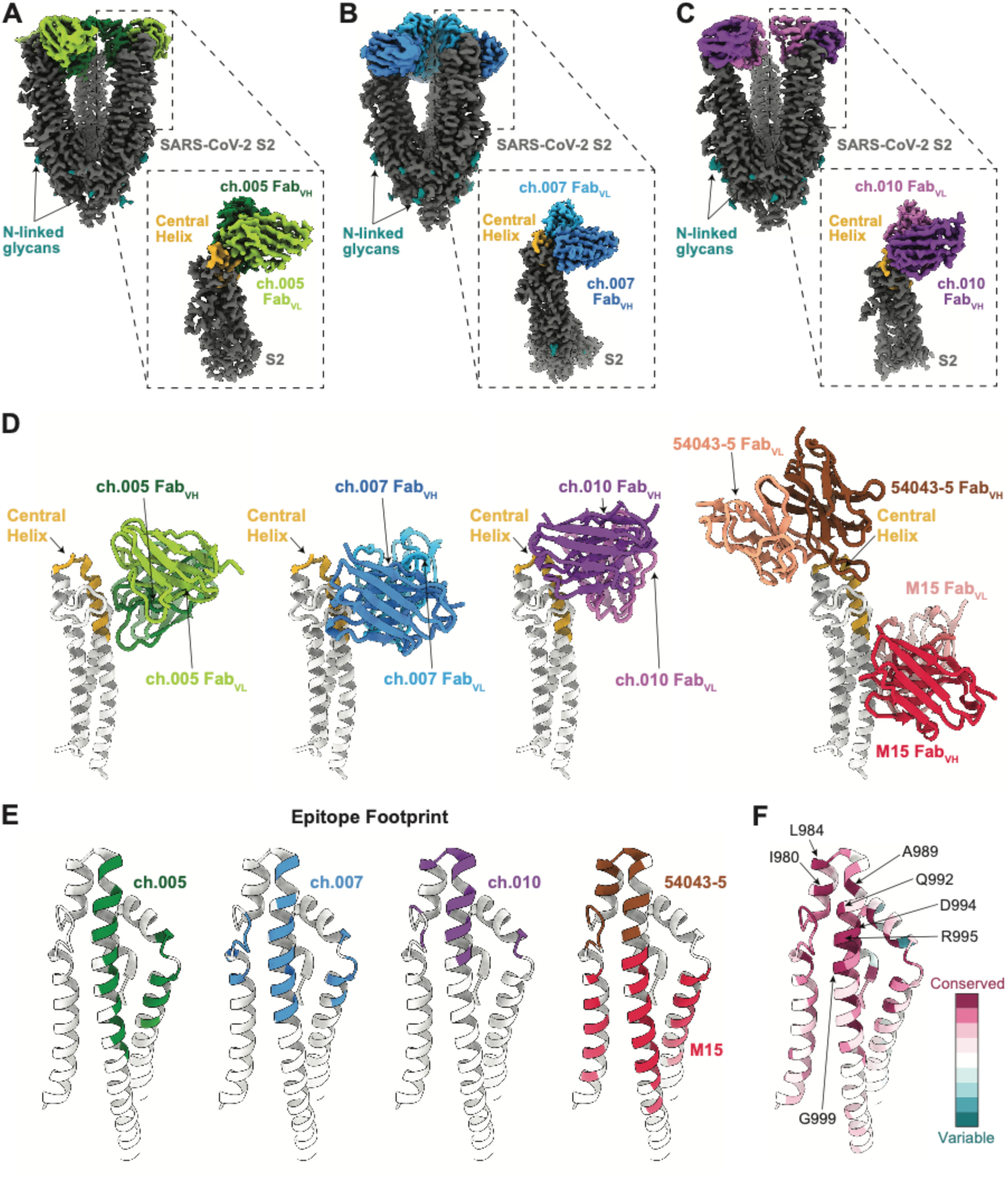
Cryo-EM of central helix antibodies complexed with S2 reveals distinct binding poses. **(A-C)** Cryo-EM density for ch.005-SS **(A)**, ch.007-SS **(B)**, and ch.010-SS **(C)** at 3.3 Å, 2.7 Å, and 3.0 Å, respectively. Insets depict the local refinement cryo-EM density for antibody V_H_ and V_L_ domains binding to a single SS protomer. Central helix colored in gold. Maps presented were produced using DeepEMhancer (*76*) through the COSMIC Cryo-EM server (*77*). **(D)** Models of antibody V_H_ and V_L_ domains binding to a single SS protomer, compared with the 54043-5 (PDB 8VCR) and M15 (PDB 9MPW) V_H_ and V_L_. SARS-CoV-2 S2 is aligned across all four models depicted to illustrate differences in binding poses. **(E)** Antibody epitope footprint depicted on the SS protomer (PDB 8VCR) as defined by buried surface area using PDBePISA (*78, 79*). **(F)** Depiction of residue conservation across 26 CoVs with representative strains (see Methods) from all four genera; produced via Consurf Server (*80*).

We next compared the CH antibody structures to the previously described 54043-5 (*37*) and M15 (*75*) antibodies (PDB 8VCR and 9MPW, respectively) (Figure 3D). Structural alignment to C*α* atoms of SS protomers (residues 736-781 and 943-1027) yielded a root mean square deviation of <0.92 Å, indicating a conserved S2 fold. Whereas 54043-5 primarily contacts the apical helical turn of the central helix through its CDRH3 loop, the antibodies reported here recognize buried residues along the inner face of the central helix. ch.005 and ch.007 targeted more distal CH residues and the neighboring helix of the same protomer, similar to M15, whereas the ch.010 footprint extended toward and partially overlapped the apical 54043-5 epitope (Figure 3D). Modeling these interactions onto the postfusion SARS-CoV-2 spike (PDB 7E9T) confirmed that their epitopes would be occluded in the postfusion state (Figure S5A), suggesting that CH antibody recognition is restricted to the prefusion conformation. Although we were unable to determine an atomic model for ch.008, docking of Fab models into the ch.008-SS local density provided insight into its binding pose, indicating that ch.008 adopts a similar orientation as ch.007 and potentially engages apical residues near the helical turn resembling ch.010 (Figure S4D). To compare epitope footprints, we calculated buried surface areas at the Fab-SS interface (Figure 3E). While all three antibodies approached the helix in a similar angle (Figure 3D), ch.005 and ch.007 targeted more distal residues, including contacts on the adjacent helix of the same protomer (residues 750-765), much like M15, whereas ch.010’s contacts overlapped with the 54043-5 footprint (Figure 3E). To better understand the basis of broad cross-reactivity, we mapped sequence conservation scores from 26 representative coronavirus strains spanning the four subgenera onto the SARS-CoV-2 S2 (Figure 3F). Notably, residues D994_SARS-CoV-2_ and R995_SARS-CoV-2_ are strongly conserved (Figure 3F), with R995_SARS-CoV-2_ forming part of the footprint for ch.005, ch.007, ch.010, and 54043-5 antibodies (Figure 3E).

Together, these structures expand the known landscape of CH recognition, revealing a continuum of antibody engagement along the conserved residues on the inner face of the central helix and providing structural insight into their broad recognition across betacoronaviruses.

### Central Helix antibodies engage conserved motifs on S2 through distinct binding modes

Given their genetic diversity and distinct binding poses, we next asked whether CH-directed antibodies converge on common molecular interactions at the Fab-spike interface (Figure 4 and S5B-E). Examination of paratope residues revealed distinct CDR loop usage among the three antibodies (Figure S5B). Both ch.005 and ch.010 relied primarily on CDRH1 and CDRH3 loops for epitope engagement, with ch.010 further stabilized by extensive framework contacts (Figure S5B). In contrast, ch.007 exhibited a paratope defined by interactions distributed among all CDR and framework regions, with the CDRH3 and CDRL3 making the most substantial contributions. Previously described antibody M15 was similarly described to utilize light chain somatic hypermutations to enhance its affinity to the CH (*75*). Examination of V gene-encoded contacts at the paratope residues showed that ch.007 had the lowest contribution of somatic hypermutations to buried surface area (16.7%), whereas ch.005 and ch.010 had higher and comparable contributions (∼27%) (Figure S5C-E). Thus, genetically distinct CH-directed antibodies use different combinations of germline-encoded and somatically mutated residues to engage this conserved region.

**Figure 4.**
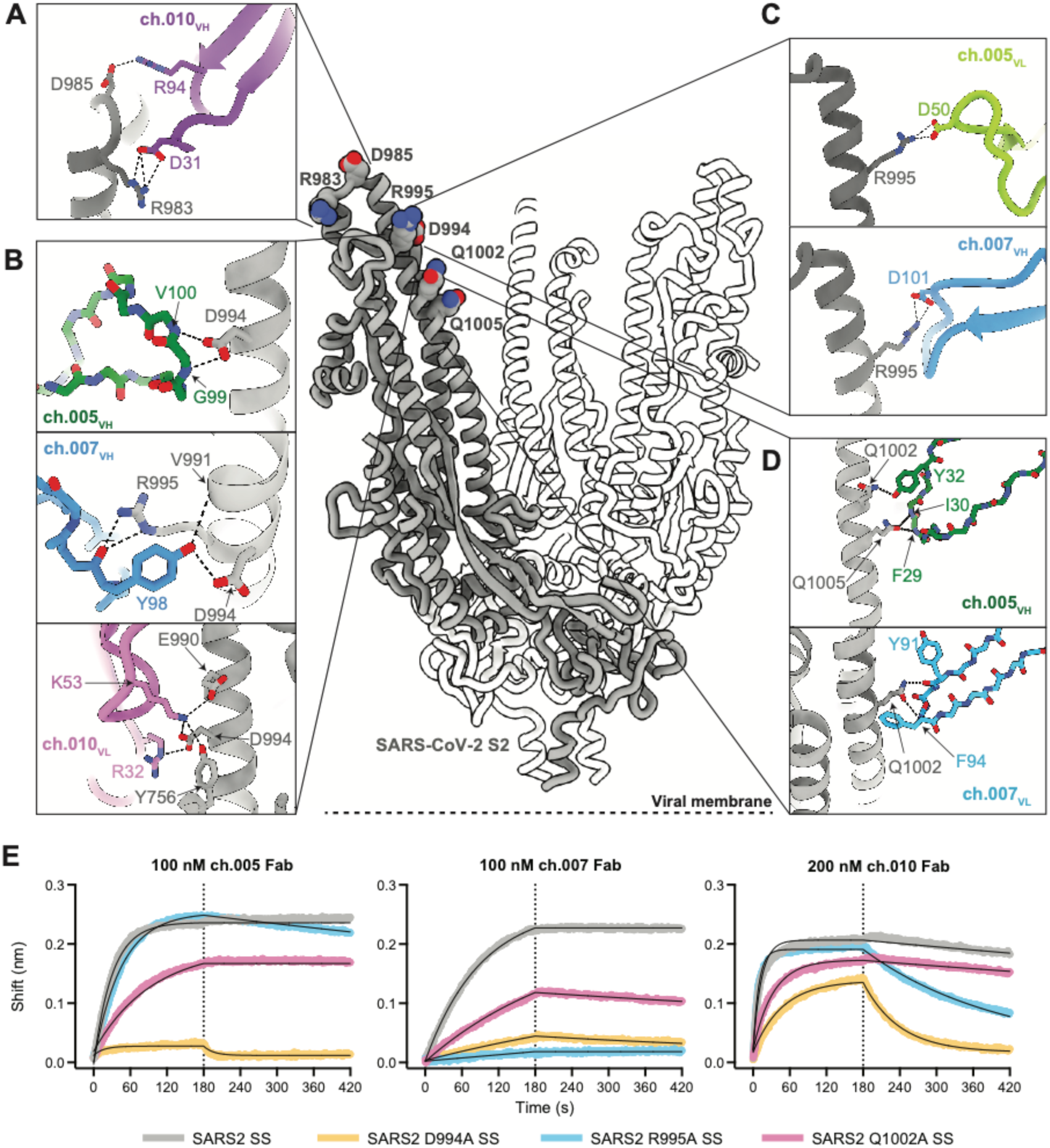
Molecular interactions between CH antibodies and the SARS-CoV-2 S2 subunit. **(A-D)** Residue-level contacts between the SS (dim gray) and ch.005, ch.007, or ch.010 Fabs. Each panel compares antibody contacts to shared epitope residues. Potential hydrogen bonds depicted by dotted black lines and defined according to PDBePISA. **(E)** Representative BLI sensorgrams of binding of monoclonal antibodies to single knock out SARS-CoV-2 epitope residues.

Despite these differences in paratope composition, examination of the specific molecular interactions across the three CH antibodies indicated convergence on highly conserved residues within the central helix (Figure 4). Our structures reveal both shared interaction hotspots and antibody-specific contacts that together explain broad recognition of this conserved epitope. Among the three, only ch.010 engaged the helical turn of the CH, forming hydrogen bonds with R983_SARS-CoV-2_ and D985_SARS-CoV-2_ (Figure 4A), interactions similar to those observed for 54043-5 (Figure 3E). All three antibodies interacted with D994_SARS-CoV-2_, though the modes of engagement differed: ch.005 via backbone hydrogen bonding, and ch.007 and ch.010 via sidechain contacts (Figure 4B). In ch.010, these interactions were further stabilized by hydrogen bonds involving K53_ch.010 VH_ and Y756_SARS-CoV-2_ and E990_SARS-CoV-2_. In addition to the substantial contacts made with D994_SARS-CoV-2_, both ch.005 and ch.007 formed potential salt bridges with R995_SARS-CoV-2_, mediated by either an aspartic acid on the light chain (ch.005) or heavy chain (ch.007) (Figure 4C). While slightly less conserved than D994_SARS-CoV-2_ and R995_SARS-CoV-2_, Q1002_SARS-CoV-2_ served as a shared contact residue for ch.005 and ch.007 (Figure 4D), with an equivalent interaction observed at Q1005_SARS-CoV-2_ in the ch.005 structure due to its slightly altered binding pose (Figure 3D-E and 4D). M15 similarly engaged Q1005_SARS-CoV-2_, forming a ternary interaction (*75*).

To experimentally validate the contribution of these shared contact residues, we introduced alanine mutations at positions D994_SARS-CoV-2_, R995_SARS-CoV-2_, and Q1002_SARS-CoV-2_ and assessed how each mutation affected antibody binding compared to the wild-type SS (Figure 4E and Table S2). Among the residues tested, substitutions at D994_SARS-CoV-2_ and R995_SARS-CoV-2_ produced the most pronounced antibody-specific effects on binding. D994A_SARS-CoV-2_ abrogated ch.005 Fab binding, while ch.010 Fab was still able to bind SS D994A_SARS-CoV-2,_ albeit with a reduced k_on_, the k_off_ rate was approximately 32 times slower compared to wild-type (Figure 4E and Table S2). While ch.005 binding was unaffected by the R995A_SARS-CoV-2_ mutation, ch.007 was unable to bind this SS variant and the k_off_ rate for ch.010 binding was approximately 8.5 times slower than the wild-type SS.

Together, these analyses reveal both conserved interaction hotspots that anchor antibody binding, particularly D994 and R995, while employing antibody-specific adaptations that diversify recognition of the central helix. This combination of convergent epitope targeting and flexible recognition strategies likely contributes to the broad coronavirus reactivity observed for CH-directed antibodies.

### Central Helix antibody ch.007 requires Fcγ receptor engagement for protection against SARS-CoV-2 challenge in vivo

To evaluate whether CH- and LS-directed antibodies could mediate antiviral functions beyond direct virus neutralization, we assessed their ability to induce antibody-dependent cellular cytotoxicity (ADCC) and antibody-dependent cellular phagocytosis (ADCP), two Fc-mediated effector functions (Figure S6A-S6B). CH-directed antibodies demonstrated no detectable ADCC or ADCP activity under the conditions tested. In contrast, several LS-directed antibodies, including ls.015, ls.017, ls.018, ls.019, and ls.020, exhibited measurable ADCC activity, with ls.017 also mediating ADCP. To determine whether these effects were Fc*γ* receptor-dependent, we generated Fc-silenced (G236R/L328R; GRLR) and Fc-enhanced (G236A/A330L/I332E; GAALIE) variants of ls.019. The observed ADCC activity was abrogated by GRLR mutations and enhanced by GAALIE mutations, confirming that the observed activity was Fc*γ* receptor dependent (Figure S6C-S6D).

We next asked whether CH- and LS-directed antibodies could confer protection *in vivo* despite their limited virus neutralizing activity. Representative CH antibodies (ch.005, ch.007, and ch.010) and the LS antibody ls.019 were first evaluated in a pilot prophylactic SARS-CoV-2 challenge study (Figure 5A). BALB/c mice (n=3 per group) received a single dose (500 µg) of antibody prior to challenge with the mouse-adapted SARS-CoV-2 MA10 isolate (*81*) and were monitored for disease-associated weight loss over a 10-day period. Among the antibodies tested, ch.007 exhibited the greatest protection, with substantially reduced weight loss relative to animals receiving an isotype control antibody (Figure 5A). Antibody ls.019 showed a more modest protective trend, prompting a higher-powered follow-up study (n=6 per group) comparing parental ls.019 with its Fc-silenced GRLR variant (Figure S7). In this experiment, neither parental ls.019 nor ls.019 GRLR exhibited significant protection from SARS-CoV-2 challenge compared to isotype control in weight loss (Figure S7A), survival trajectories (Figure S7B), or infectious virus titers in the lungs (Figure S7C), and no significant differences were observed between the WT and GRLR variant.

**Figure 5.**
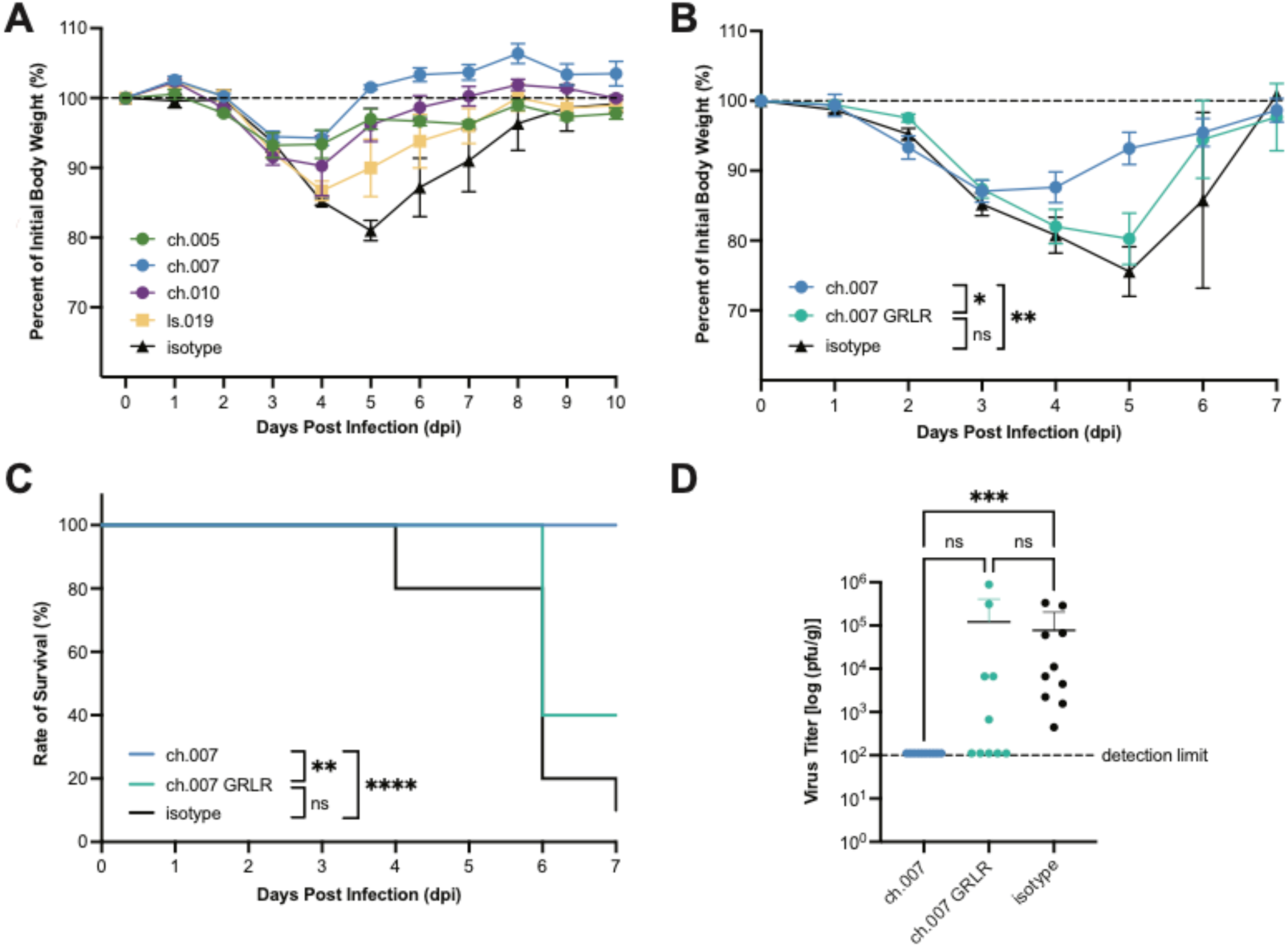
Protection by central helix antibodies in a SARS-CoV-2 MA10 mouse model. **(A-B)** Body weight curves for BALB/c mice treated prophylactically with 500 µg of either (A) the indicated central helix and lower stalk antibodies or (B) ch.007 and ch.007 with G236R/L328R mutations (ch.007 GRLR), compared to mice treated with isotype control antibodies (500 µg), prior to intranasal (i.n.) inoculation with the mouse-adapted SARS-CoV-2 MA10 strain. Panel (A) represents a pilot experiment with n = 3 per group; error bars represent the standard error of the mean but statistical analysis was not performed. For panel (B), error bars represent the standard error of the mean of the mice within each test group (n = 10). Statistical significance was determined at 5 dpi by one-way ANOVA followed by Tukey’s multiple comparisons test. *p < 0.05, **p < 0.01; ns, not significant. **(C)** Survival curves for mice treated with ch.007 compared to ch.007 GRLR and isotype control from panel (B). Statistical significance was logrank (Mantel-Cox) test **p < 0.01, ***p < 0.001; ns, not significant. **(D)** Quantification of viral titer within lungs of mice treated with ch.007, ch.007 GRLR, or isotype control from panel (B). Statistical significance was determined by Kruskal-Wallis test followed by Dunn’s multiple comparisons test. *p < 0.05, **p < 0.01, ***p < 0.001; ns, not significant.

Despite the absence of detactable ADCC or ADCP activity *in vitro* (Figure S6A-S6B), and given the protection observed with ch.007 in the pilot study, we compared parental ch.007 against the ch.007 GRLR variant in a parallel prophylactic challenge experiment (Figure 5B-D, n=10 per group). Mice treated with the ch.007 GRLR variant exhibited markedly diminished protection, with weight loss trajectories (Figure 5B), survival rates (Figure 5C), and lung viral titers (Figure 5D) mirroring those of animals treated with the isotype control antibody. In contrast, all mice treated with ch.007 survived through the study endpoint (Figure 5C) and no infectious virus was detected in their lungs (Figure 5D).

Together, these data demonstrate that Fc*γ* receptor engagement is required for ch.007-mediated protection *in vivo*, despite the absence of detectable ADCC or ADCP activity in the assays tested. Conversely, the Fc*γ* receptor-dependent ADCC activity observed for LS antibodies *in vitro* did not translate into a measurable Fc-depedent protective effect for ls.019 *in vivo*. These findings highlight that individual Fc effector assays do not necessarily predict the protective activity of S2-directed antibodies and further underscore that the antiviral potential of CH-directed antibodies is not adequately predicted by virus neutralizing or antibody effector activity alone.

## DISCUSSION

The persistent threat of zoonotic coronavirus emergence underscores the need to identify conserved and functionally constrained regions on the spike glycoprotein that can be targeted for broadly protective immunity. While much of the coronavirus antibody response targets the receptor binding domain (RBD) within S1, the more conserved S2 domain has recently emerged as an important source of cross-reactive antibody epitopes to inform pan-coronavirus vaccine and antibody strategies (*16*). Here, we define genetically and structurally distinct classes of S2-directed human antibodies that engage the central helix (CH) and the membrane-proximal lower stalk (LS) of spike. Our findings expand the known landscape of human antibody recognition across the conserved S2 fusion protein and reveal that CH-directed responses encompass greater genetic, structural, and functional diversity than previously appreciated.

We previously used evolutionary sequence conservation to identify antibody-accessible “coldspots” within spike and demonstrated that the FP and HR2 stem helix (SH) are targeted by broadly reactive human antibodies with virus neutralizing and protective activity (*23*). Here, analysis of more than 17 million SARS-CoV-2 sequences collected over several additional years of viral evolution demonstrated that these S2 coldspots remain remarkably stable, while also identifying a conserved region within the membrane-proximal LS. Although antibody responses to the iFP, CD, CH and LS were less frequently detected than those targeting the FP and SH coldspots, we isolated human monoclonal antibodies against each region (Figure 1). In contrast to the previously characterized FP and SH monoclonal antibodies (*23*), antibodies targeting these additional coldspots in S2 exhibited minimal neutralizing activity. However, CH- and LS-directed antibodies recognized highly conserved epitopes and displayed broad reactivity across SARS-CoV-2 variants and divergent CoVs, including zoonotic CoVs with spillover potential. These findings demonstrate that evolutionary conservation across S2 supports antibody responses with distinct functional properties and extend the coldspot framework beyond conserved sites associated with direct virus neutralization.

We describe and characterize a LS-directed antibody ls.019 which targets a membrane-proximal region of S2 encompassing SARS-CoV-2 residues 1177–1190. This region overlaps with the HR2 domain, which forms part of the six-helix bundle in the postfusion state and is essential for membrane fusion. Although antibodies and nanobodies recognizing overlapping HR2 regions have been described, the ls.019-peptide structure provides, to our knowledge, the first high-resolution molecular view of a human-derived antibody engaging this conserved LS epitope. Unexpectedly, the crystal structure of the ls.019–peptide complex revealed that the LS bound peptide adopted an extended rather than the predicted prefusion helical conformation, suggesting conformational adaptability in epitope recognition by ls.019. Although the conformation of the LS epitope in the context of antibody-bound native spike remains unresolved, these findings extend the structurally defined landscape of human S2 immunity beyond the FP, CH, and canonical SH toward the membrane-proximal base of spike. Moreover, several LS-directed antibodies mediated Fc*γ* receptor-dependent ADCC and ADCP activity *in vitro*, demonstrating that antibodies recognizing this membrane-proximal S2 region can engage cellular effector pathways despite their limited neutralizing activity.

Our cryo-EM analyses of CH-directed antibodies reveal a continuum of binding poses extending from the S2 apex distally along the inner face of the CH, each engaging conserved residues critical for the prefusion-to-postfusion transition. Previous studies have identified recurrent antibody responses to the CH/S2 apex, including the public IGHV1-69/IGKV3-11 antibody class (*38, 82*), the structurally distinct antibodies 54043-5 and M15 (*37, 75*), and highly expanded public clonotypes with restricted germline usage identified through polyclonal analyses (*40*). Collectively, these studies have established the CH/S2 apex as a recurrent antibody target, with much of the previously described recognition concentrated toward apical regions and associated with convergent or restricted genetic solutions. Our structures expand this emerging landscape by demonstrating that recognition of the CH can be achieved through multiple independent genetic and structural solutions. Antibodies ch.005, ch.007, and ch.010 recognize buried residues along the inner CH through binding poses and genetic solutions distinct from those of the apically focused IGHV1-69/IGKV3-11 class and previously structurally characterized CH antibodies. Despite converging on overlapping epitope footprints, these antibodies use distinct V genes, CDR architectures, and combinations of germline-encoded and somatically mutated contacts. Thus, alongside recurrent public responses, our findings demonstrate that genetically distinct human antibody lineages can independently converge on conserved features of the S2 apex through multiple structural solutions, reflecting intrinsic plasticity in the human naïve repertoire for recognizing the conserved CH epitope.

At the molecular level, the structurally diverse CH antibodies converged on two highly conserved residues, specifically D994_SARS-CoV-2_ and R995_SARS-CoV-2_. These residues contribute to stabilization of the prefusion trimeric core, with previous structural studies implicating them in regulating S2 refolding during membrane fusion (*73*). Their conservation, therefore, likely reflects functional constraints that create durable sites for antibody recognition, although mutations within or surrounding these epitopes may still alter antibody engagement through direct or conformational mechanisms. Our findings support the emerging view that the S2 subunit, long thought to be poorly accessible, harbors multiple conformationally accessible epitopes that can be recognized by antibodies. Although binding to this region could influence S2 conformational dynamics, the limited neutralizing activity of CH-directed antibodies suggests that direct inhibition of membrane fusion is unlikely to explain their antiviral effects.

Indeed, an important finding of this study is that the poorly neutralizing but broadly reactive CH antibody ch.007 conferred protection in a SARS-CoV-2 MA10 challenge model. Protection was lost following introduction of Fc-silencing GRLR mutations, establishing that Fc*γ* receptor engagement is mechanistically required for ch.007-mediated protection *in vivo*. Fc-dependent protection by non-neutralizing antibodies targeting the S2 core has also been reported for members of the IGHV1-69/IGKV3-11 public antibody class. However, this protective phenotype does not appear to be shared uniformly across CH- and S2 apex-directed antibodies, which are predominantly weakly or non-neutralizing, and have shown limited or no protective activity (*37, 39, 40*). Recent polyclonal analyses have further concluded that immunodominant responses to portions of the S2 apex may represent broadly reactive but functionally limited antibody responses (*40*). Our findings extend Fc-dependent protection to a genetically and structurally distinct mode of CH recognition, indicating that protective activity is not restricted to a recurrent public antibody class. Rather, the divergent protective phenotypes reported across neighboring S2 epitopes suggest that CH- and S2 apex-directed antibodies can access distinct pathways to antiviral protection.

Unexpectedly, however, ch.007 and the other CH-directed antibodies exhibited no detectable ADCC or ADCP activity in the assays tested here, despite the requirement for Fcγ receptor engagement in ch.007-mediated protection *in vivo*. Conversely, multiple LS-directed antibodies mediated robust Fc*γ* receptor-dependent activity *in vitro*, yet Fc silencing of ls.019 did not significantly alter protection *in vivo*. Thus, across these two conserved S2 epitopes, measurable Fc effector activity *in vitro* did not predict Fc-dependent protection *in vivo*. Similar discordance has been observed for other antiviral antibodies. The S2 apex-directed antibody 54043-5 mediated ADCP and antibody-dependent cellular trogocytosis *in vitro* but failed to protect against SARS-CoV-2 challenge (*37*). Likewise, although ADCC generally correlated with protection among non-neutralizing influenza HA-directed antibodies, some individual antibodies protected despite little or no detectable ADCC, whereas another with substantial ADCC activity failed to protect *in vivo* (*83*). Together with our findings, these observations suggest that individual *in vitro* Fc effector assays may not reliably predict antibody protection and likely capture only a subset of the mechanisms contributing to Fc-dependent antiviral activity *in vivo*.

The consequences of Fc*γ* receptor engagement also appear to vary among non-neutralizing S2-directed antibodies. Similar to ch.007, Fc-dependent protection has been demonstrated for the non-neutralizing IGHV1-69/IGKV3-11 S2-apex antibody COVA2-18, for which Fc attenuation abrogated protection against lethal SARS-CoV-2 challenge (*84*). In contrast, wild-type 54043-5 failed to protect *in vivo,* whereas partial protection emerged following introduction of the Fc-attenuating LALA-PG substitutions (*37*). Although differences between experimental models preclude direct mechanistic comparisons, these divergent phenotypes further indicate that S2-directed antibodies cannot be considered a functionally uniform class. Epitope position, binding geometry, antigen accessibility, immune-complex architecture, and recognition of spike on infected cells may each influence the consequences of Fc*γ* receptor engagement and ultimately determine protective activity. Furthermore, other antibody-dependent antiviral mechanisms may also extend beyond the cellular effector pathways examined here. Recent work demonstrated that complement can enhance the antiviral activity of cross-reactive S2 antibodies, including antibodies with otherwise limited neutralizing activity through C1q and C3-dependent mechanisms (*85*), further illustrating that ADCC and ADCP assays may not capture the full range of antibody effector mechanisms operating *in vivo*. Together, these observations highlight the diversity of effector mechanisms available to S2-directed antibodies and the importantance of antibody-specific properties in determining their potential for vaccines and antibody-based interventions.

Several limitations should be considered when interpreting these findings. Although Fc-silencing established a requirement for Fcγ receptor engagement in ch.007-mediated protection *in vivo*, the specific Fc receptors, effector cells, and antiviral mechanisms responsible remain unresolved. The absence of detectable ADCC or ADCP activity in our *in vitro* assays further emphasizes this mechanistic gap. Conversely, although LS-directed antibodies mediated Fc*γ* receptor-dependent activity *in vitro*, ls.019 did not confer significant protection *in vivo*, indicating that these measureable Fc effector activities were insufficient to translate into protection in this model. More broadly, protection was evaluated only against SARS-CoV-2 MA10, and whether the broad binding activity of CH-directed antibodies translates into protection against divergent coronaviruses remains unknown. Structurally, limited local resolution prevented atomic modeling and complete characterization of the mechanism of binding of ch.008 to the CH, restricting our interpretations only to general binding pose as shown in the supplemental figure. Similarly, although the ls.019 Fab-peptide structure defines the molecular determinants of LS recognition, the conformation and accessibility of this epitope when engaged within native prefusion spike remain unresolved.

Together, our findings expand the landscape of conserved antibody vulnerabilities within the coronavirus S2 subunit and reveal greater genetic, structural and functional diversity among S2-directed antibody responses than previously appreciated. Genetically distinct antibodies can converge on conserved features of the CH through multiple binding solutions, while human antibody recognition extends farther toward the viral membrane into the HR2 lower stalk. Importantly, the discordance between neutralization, Fc effector activities measured *in vitro*, and protection *in vivo* demonstrates that the antiviral potential of antibodies targeting conserved S2 epitopes cannot be defined by any single functional measurement. Rather, our findings support a broader view of S2 immunity in which conserved epitopes can support diverse antibody responses whose protective potential emerges from the combination of epitope recognition, binding geometry and engagement of distinct antiviral effector mechanisms. Defining and selectively eliciting these protective responses may provide new opportunities for the development of broadly protective coronavirus vaccines and antibody-based countermeasures.

## MATERIALS AND METHODS

### Human subjects

Plasma and peripheral blood mononuclear cells (PBMC) samples used in the experiments reported in Figure 1 and S1 were obtained from COVID-19 convalescent study participants enrolled in the Lugano cohort (Clinica Luganese Moncucco, Switzerland) (*48*). The individuals were diagnosed with COVID-19 in the period March to November 2020, and the samples obtained 83-269 days from symptoms onset(*48*). Control plasma samples were from individuals with no prior SARS-CoV-2 infection or vaccination, as confirmed by negative serologic test. The study was performed in compliance with all relevant ethical regulations and study protocols approved by the Ethical Committee of the Canton Ticino: CE-3428 and CE-3960.

### Cell lines

Human embryonic kidney (HEK) 293T cells (*Homo sapiens*, embryonic kidney cells) were used for pseudovirus generation and cultured in Dulbecco’s Modified Eagle Medium (DMEM) supplemented with 10% fetal bovine serum (FBS). HEK293T cells overexpressing human angiotensin-converting enzyme 2 (ACE2; HEK293T_ACE2_) used for pseudovirus titration and neutralization assays were generated as previously described (*23*) and were cultured in DMEM supplemented with 10% FBS, 1% non-essential amino acids, 1 mM sodium pyruvate, 1x penicillin/streptomycin, and 5 µg mL^−1^ blasticidin. HEK293T cells used for detecting antibody binding to cell-expressed spike by flow cytometry were co-transfected with plasmids encoding for green fluorescent protein (GFP) and the spike protein in a 1:1 ratio using PEI-MAX (Polysciences) as previously described (*23*). Expi293F™ cells (Thermo Fisher Scientific, cat. no. A14527) were used for recombinant expression of monoclonal IgGs and viral antigens and were cultured in Expi293 expression medium (Thermo Fisher Scientific, cat. no. A1435014) according to the vendor’s instructions.

### Blood samples processing and storage

Peripheral blood mononuclear cells (PBMCs) were isolated by Histopaque density centrifugation and stored in liquid nitrogen in the presence of FBS and DMSO. Plasma was aliquoted and stored at −20°C or less. Prior to experiments, aliquots of plasma were heat-inactivated (56°C for 1 hour) and then stored at 4°C.

### Computational analyses of coldspot sequences

Analysis of SARS-CoV-2 sequences and subsequent coldspot determination (Figure S1A and Table S1) were performed using the coldspot data pipeline available on GitHub (https://github.com/cavallilab/coldspot) as previously described (*23*), except that sequences of spike that were present in the GISAID database up to July 23, 2025 and had a length of 1223-1323 aa and no undetermined aa were included.

### Peptides

Peptides used for binding assays, sorting by flow cytometry, and structural studies were synthesized by Genscript as previously described (*23*). In brief, all peptides were synthesized with a biotin-amino hexanoic acid (Ahx) and amide at the N- and C-terminus, respectively. Peptides were resuspended in 100% dimethyl sulfoxide (DMSO) and stored at −20°C. Sequences of all peptides used in this study are described in Table S1.

### Single B cell sorting

B cells were sorted from PBMCs as previously described (*23*). In brief, B lymphocytes were enriched (Miltenyi Biotec, 130–101-638) prior to labeling with anti-CD20-PE-Cy7 (BD Biosciences, 335828), anti-CD14-APC-eFluor 780 (Thermo Fischer Scientific, 47–0149-42), anti-CD16-APC-eFluor780 (Thermo Fischer Scientific, 47–0168-41), anti-CD3-APC-eFluor 780 (Thermo Fischer Scientific, 47–0037-41), anti-CD8-APC-eFluor 780 (Invitrogen, 47–0086-42), Zombie NIR (BioLegend, 423105), as well as fluorophore-labeled ovalbumin (Ova) and peptides. B cells labeling as ZombieNIR^−^CD14^−^CD16^−^CD3^−^CD8^−^CD20^+^Ova^−^peptide-PE^+^peptide-AF647^+^ were sorted as single cells using a FACSymphony S6 (Becton Dickinson). The sorting strategy is shown in Figure S1C.

### Antibody gene sequencing, cloning and expression for ELISA, neutralization assays, and in vivo experiments

The identification of antibody gene sequences was as described previously (*86*). In brief, RNA from single cells was reverse-transcribed (SuperScript III Reverse Transcriptase, Invitrogen, 18080–044) prior to nested PCR amplification of the variable IGH, IGL and IGK genes and Sanger sequencing and cloning into IgG1 expression vectors. Recombinant monoclonal antibodies used in Figures 1, 5, and S2 were produced by transient transfection of Expi293 cells and purified from supernatants, as previously described (*87*).

### Antibody production for crossreactivity ELISA and BLI

Monoclonal IgGs were expressed as previously described (*88*). In brief, plasmids encoding the heavy IgG and LCs for each antibody were co-transfected in Expi293F cells using the ExpiFectamine transfection kit (Thermo Fisher Scientific, cat. no. A14525) in a 1:1 ratio. ExpiFectamine transfection enhancers were added 20 hr post transfection, per manufacturer’s protocol. Cell supernatants were harvested, filtered, and subsequently ran over HiTrap MabSelect SuRe columns (Cytiva) to capture expressed IgGs. Elutions were concentrated using Amicon ultra-15 centrifugal filters with a 30 kDa molecular weight cut off (MWCO) (Millipore, cat. no. UFC9030) and purified via size exclusion chromatography (SEC) using a HiLoad 16/600 Superdex 200 pg or Superdex 200 Increase 10/300 GL column (Cytiva) into 1x Tris-buffered saline supplemented with sodium azide. (TBS-Az: 20 mM Tris-HCl pH 8.0, 150 mM sodium chloride, 0.02% (v/v) sodium azide).

### Antibody production for structural studies

Antibody Fabs used for X-ray crystallography and cryo-EM were prepared via papain cleavage of IgGs expressed, as described above. In brief, papain (Sigma-Aldrich, cat. no. P3125) was incubated for 10 min in 2x Digestion Buffer (50 mM NaPO4 pH 7.0, 20 mM ethylenediaminetetraacetic acid (EDTA)) at 37°C at a final mass 1.5% (w/w) of the total IgG protein to be cleaved. Activated papain was then added to the IgGs at a 1:1 ratio (v/v) and incubated at 37°C for 1 hr with shaking at ∼100 rpm. The enzymatic reaction was quenched with iodoacetamide (Sigma-Aldrich, cat. no. I1149) and then ran over a HiTrap MabSelect SuRe column (Cytiva) to separate uncleaved IgG and cleaved Fc from the desired Fab product. Cleaved Fabs were further purified via SEC on a HiLoad 16/600 Superdex 200 pg or Superdex 200 Increase 10/300 GL column (Cytiva) using 1x TBS-Az and concentrated using Amicon ultra-15 centrifugal filters with 10 kDa MWCOs (Millipore, cat. no. UFC9010).

The ls.019 antibody single chain variable fragment (scFv; V_H_-(G_4_S)_4_-V_L_-GS-His_6_) gene was ordered from Integrated DNA Technologies (IDT) and cloned into the AbVec2.1 expression plasmid using Gibson assembly. The scFv was expressed in Expi293F cells, as described for IgGs above, and subsequently purified from cellular supernatants via affinity chromatography using a HisTrap column (Cytiva). Eluted scFv was further purified via SEC on a Superdex 200 Increase 10/300 GL column (Cytiva) into 1x TBS using an AKTA pure M1 system (Cytiva).

### Recombinant spike protein production

Sequences for the recombinant spike proteins were codon optimized and cloned into a mammalian expression vector with an N-terminal secretory signal peptide and a variation of C-terminal tags including the T4 Foldon trimerization domain, polyhistidine tag for affinity purification, and biotinylation motif. The SARS-CoV-2 SS construct used for binding and structural studies was designed as a combination of the previously described HexaPro-S2 and HexaPro-SS designs (*72*). In brief, the construct encoded for residues 696-1213_SARS-CoV-2_ with the following stabilizing mutations: T696Q, S704C/K790C, Q957E, and the HexaPro mutations (F817P, A892P, A899P, A942P, K986P, and V987P). Plasmids were transfected in Expi293F following manufacturer instructions and using the Expi293 Expression System Kit (Gibco, cat. no. A14635). To obtain biotinylated proteins, Expi293F cells were co-transfected with a 1:1 ratio of the spike construct DNA and a plasmid encoding for the *Escherichia coli* biotin ligase (BirA) enzyme. Recombinant proteins were subsequently isolated from cellular supernatants via affinity chromatography using a HisTrap column (Cytiva). Further purification was performed via SEC on either a HiLoad 16/600 Superose 6 pg or Superose 6 Increase 10/300 column (Cytiva) using an AKTA pure M1 system (Cytiva).

### Enzyme-linked immunosorbent assays (ELISAs)

#### ELISAs to measure the binding of plasma IgGs and human monoclonal IgGs to coldspot peptides

ELISAs to screen for patient plasma IgG reactivity to coldspot peptides (Figure 1A, 1C, and S1B) and subsequent quantification of binding of recombinantly expressed monoclonal IgGs to coldspot peptides (Figure S2A-B) were performed as previously described (*23*). In brief, 384-well plates (Thermo Fisher Scientific, cat. no. 464718) were prepared by incubating biotinylated peptide resuspended in phosphate-buffered saline (PBS) for 1 hr at room temperature in wells pre-coated with NeutrAvidin (2 µg mL^−1^) overnight at room temperature. Following coating, plates were blocked for 2 hr at room temperature using PBS + 2% bovine serum albumin + 0.05% Tween 20. After a wash step, serial dilutions of plasma samples (assayed at a starting dilution of 1:50 and subsequently diluted three-fold) or monoclonal antibodies were incubated with the immobilized peptide for 1 hr at room temperature to identify anti-coldspot IgGs. Following the incubation and a wash step, plates were incubated with anti-human IgG secondary antibody conjugated to horseradish peroxidase (HRP) (GE Healtchare, NA933) at a 1:5000 dilution. Last, plates were developed by the addition of HRP substrate 3,3’,5,5’-tetramethylbenzidine (Thermo Fisher Scientific, cat. no. 34021) for 10 min and then quenched with 1 M H_2_SO_4_ solution. Absorbance was measured at 450 nm using a microplate reader (BioTek) equipped with Gen5 software.

#### ELISAs to measure the binding of human monoclonal antibodies to SARS-CoV-2 spike protein

ELISAs to evaluate binding of recombinantly-expressed monoclonal IgGs to the SARS-CoV-2 spike-2P stabilized ectodomain (Figure 1E and S2A) were performed in a similar manner as for the coldspot peptides, described above, with minor differences. Spike was coated onto 384-well plates at a concentration of 5 µg mL^−1^ in PBS overnight at room temperature. Following blocking and washing, monoclonal IgGs were tested in serial dilution. Readout and data analysis were the same as described above.

#### ELISAs to measure the binding of human monoclonal antibodies to CoV spike proteins

ELISAs to measure the binding crossreactivity of monoclonal IgGs to CoV spike proteins (Figure 2) were performed as previously described (*88*). In brief, 200 ng of the recombinant spike proteins were immobilized directly on high-binding 96-well assay plates (Corning, cat. no 9018) in 1x TBS-Az for 20 hr at 4°C. Plates were then washed with ELISA washing buffer (TBS-T: 1x TBS + 0.05% Tween-20) and blocked for 1 hr at room temperature in ELISA blocking buffer (TBS-TMS: TBS + 0.05% Tween-20 + 1% non-fat dry milk + 1% goat serum). Monoclonal IgGs were incubated with immobilized spike proteins at 10-fold serial dilutions at a top concentration of 75 µg mL^−1^ in TBS-TMS buffer. Following a wash with TBS-T, HRP-conjugated goat anti-human Ig Fc antibody (Southern Biotech, cat. no. 2047-050) was used as a secondary at a 1:4000 final dilution in TBS-TMS. Following another wash with TBS-T, plates were read out with 1-Step Ultra TMB-ELISA substrate (Thermo Scientific, cat. no. 34029), quenched with 1 M HCl solution, and absorbance was measured at 450 nm with an Infinity M Plex microplate reader (Tecan) using iControl 2.0 software. Data values obtained were subtracted by the background control wells (no antigen or primary antibody added) to calculate OD_corrected_ values. EC_50_ and pEC_50_ were calculated using the [Agonist] vs. response --- Variable slop (four parameters) model in GraphPad Prism v10. EC_50_ values were only reported if three conditions were met: (1) non-linear regression fit followed a sigmoidal curve and OD_corrected_ values were both (2) greater than five times the standard deviation of the background-only wells and (3) greater than 0.5 for at least two consecutive dilutions.

### Biolayer interferometry

BLI assays were performed on an Octet RED96 system (FortéBio, Sartorius) at room temperature (20-25°C) in Octet Buffer: TBS supplemented with 0.1% (w/v) bovine serum albumin (Roche, cat. no. 03116956001) and 0.02% (v/v) Tween-20 (Sigma-Aldrich, cat. no. P9416). For assays used to calculate binding kinetics, CoV spike, SS trimers, or peptides were immobilized on streptavidin (SA) biosensors (Sartorius, cat. no. 18-5019) and dipped into a concentration series of monomeric Fabs. After subtraction of reference signal, the *k*_a_ (on rate), *k*_d_ (off rate), and dissociation constant (K_d_) were derived using the Association then Dissociation module in GraphPad Prism v10.

### Central Helix sequence conservation analysis

Full-length spike sequences from the four CoV genera (n = 26) were aligned and assessed for sequence conservation using the ConSurf server(*80*). Sequences used for the conservation analysis in Figure 2 were as follows: BkCoV-HKU9 (UniProt A3EXG6), HKU11-934 (UniProt 6BVDW0), PCoV-GX-P2V (UniProt A0A6G9KP06), HKU15 (UniProt H9B0X8), BwCoV (UniProt B2BW33), rsSHC014 (UniProt U5WLK5), BM48-31 (NCBI 014470.1), Banal 20-236 (Genbank MZ937003.2), PDF-2180 (NCBI 034440.1), WIV-1 (Genbank KF367457.1), Pangolin CoV GX_P5L (UniProt A0A6G6A1M4), rs4084 (UniProt A0A2D1PX29), BtCoV HKU4 (UniProt A3EX94), BtCoV HKU2 (UniProt A8JP00), BCRP3-Rp3 (UniProt Q3I5J5), BCHK3-HKU3 (UniProt Q3LZXI), RaTG13 (Genbank MN996532.2), MERS-CoV (UniProt K9N508), hCoV-HKU1 (UniProt Q5MQD0), hCoV-OC43 (UniProt P36334), hCoV-NL63 (UniProt Q6Q1S2), hCoV-229E (UniProt P15423), SARS-CoV (UniProt P59594), SARS-CoV-2 (UniProt P0DTC2), PdCoV (UniProt W809Y7), IBV (UniProt P12651).

### Detection of monoclonal antibody binding to spike by flow cytometry

Evaluation of IgG binding to spike presented on the cell surface was performed using flow cytometry as previously described (*23*). In brief, monoclonal IgGs were fluorescently labeled using the DY-647P1-NHS-ester reagent (Dyomics, cat. no. 647P1-01) according to the vendor’s instructions. Forty hours post generation of GFP^+^S^+^ HEK293T cells (see Cell Lines subsection), cells were reacted with monoclonal IgGs (10 µg mL^−1^) in the presence or absence of human soluble ACE2 (30 µg mL^−1^) in PBS supplemented with 5% FBS + 2 mM EDTA for 2 hr at room temperature (*23*). Following the incubation, cells were washed and then run on a FACSCanto flow cytometer (BD Biosciences); data were analyzed with FlowJo software.

### SARS-CoV-2 pseudotyped reporter viruses

The generation of spike-pseudotyped reporter viruses was performed as previously described (*89*). In brief, plasmids encoding for a C-terminally truncated spike protein (pSARS-CoV-2-S_trunc_) were co-transfected with pHIV_NL_GagPol and pCCNanoLuc2AEGFP plasmids in HEK293T cells using PEI-MAX (Polysciences). Supernatant from transfected cells was harvested at 24 hr post transfection, filtered, stored at −80°C and subsequently titrated on HEK293T_ACE2_ cells.

### Pseudotyped virus neutralization assay

Neutralization assays were performed as previously (*89*). In brief, serially diluted monoclonal antibodies were incubated with the SARS-CoV-2 pseudotyped virus for 1 hour at 37°C degrees prior to adding to HEK293T_ACE2_ cells for 48 hours. Upon washing with PBS once, cells were lysed with Luciferase Cell Culture Lysis 5x reagent (Promega) and Nanoluc Luciferase activity of lysates measured using the Nano-Glo Luciferase Assay System with GloMax Discover System reader (Promega). Luminescence units were relative to those derived from cells infected with SARS-CoV-2 pseudotyped virus in the absence of monoclonal antibodies. The half-maximal inhibitory concentration of monoclonal antibodies (IC_50_) was determined using four-parameter nonlinear regression curve fit (GraphPad Prism).

### Reporter assays for Fcγ receptor activation (ADCC and ADCP)

Fcγ receptor (FcγR) activation reporter assays were adapted from a previously reported protocol (*90*), using the optimized assay conditions described therein with minor modifications. Antibodies were diluted in Dulbecco’s modified Eagle’s medium (DMEM). SARS-CoV-2 Spike–expressing HEK293 cells (293-SARS2-S; InvivoGen, Cat. no. 293-cov2-s), maintained in DMEM supplemented with 10% FBS and penicillin-streptomycin according to the manufacturer’s instructions, were used as target cells. Diluted antibodies (20 μL) were incubated with 90 μL of 2 × 10^5^ target cells per well in 96-well plates for 1 hr at 37°C and 5% CO₂.

Jurkat-Lucia NFAT-CD16 cells expressing human CD16A/FcγRIIIA (V158 allotype; InvivoGen, Cat. no. jktl-nfat-cd16) or Jurkat-Lucia NFAT-CD32 cells expressing human CD32A/FcγRIIA (H131 allotype; InvivoGen, Cat. no. jktl-nfat-cd32) were maintained in Iscove’s modified Dulbecco’s medium (IMDM) supplemented with 10% heat-inactivated fetal bovine serum (FBS) and penicillin-streptomycin according to the manufacturer’s instructions. 90 μL of reporter cells (2 × 10^5^ cells per well) were added to the target cell–antibody mixtures, and cocultures were incubated for an additional 18 hr at 37°C and 5% CO₂. Jurkat-Lucia NFAT-CD16 and Jurkat-Lucia NFAT-CD32 cells were used to assess FcγRIIIA- and FcγRIIA-dependent activation, respectively, as surrogate readouts of ADCC- and ADCP-associated Fc effector activity.

Following incubation, 20 μL of culture supernatant was transferred to a 96-well black plate and mixed with 50 μl of QUANTI-Luc 4 Lucia/Gaussia reagent (InvivoGen, Cat. no. rep-qlc4lg1) according to the manufacturer’s instructions. Luminescence was measured immediately using a GloMax Discover Microplate Reader (Promega).

For single-concentration screening, antibodies were tested at 10 μg/mL. Fcγ receptor–dependent reporter activity was expressed as fold induction relative to the corresponding isotype control and was calculated by dividing the relative light units (RLU) value of each test condition by the RLU value of wells containing target cells, reporter cells, and the corresponding isotype control antibody, W007 (*91*). For dose-response experiments, antibodies were tested in consecutive five-fold serial dilutions, starting from 10 μg/mL. Dose-response measurements were performed in duplicate in two independent experiments. Raw RLU values and Fcγ receptor–dependent reporter fold induction were analyzed and plotted using GraphPad Prism version 10.6.1 (GraphPad Software).

### In vivo protection experiments

Mouse experiments were performed in accordance with Czech laws and guidelines on the use of experimental animals (Animal Welfare Act No. 246/1992 Coll.) and according to relevant protocols approved by the Ethics Committee of the Institute of Parasitology, Biology Centre of the Czech Academy of Sciences, and by the Departmental Expert Committee for Approval of Projects of Experiments on Animals of the Czech Academy of Sciences (approval 122/2025). SARS-CoV-2 MA10, mouse-adapted variant based on the USA-WA1/2020 backbone, was obtained through BEI Resources. The virus was used in this study after propagation on Vero E6 cells. Six-week-old female BALB/cOlaHsd mice (Envigo) were infected intranasally with SARS-CoV-2 MA10 (1 × 10^3^ plaque-forming units); in a total volume of 30 μl DMEM. Twenty-four hours before (pre-exposure prophylaxis) infection, mice were injected intraperitoneally with anti-SARS-CoV-2 monoclonal antibodies or isotype controls at the indicated amounts. Isotype control antibodies were Z021 (*87*) for Figure S6E and W014 (*91*) for Figure 5A-5C. The weight of the mice was monitored over time and at culling their tissues were collected for analysis. Lung tissues were mechanically homogenized using a Mixer Mill MM400 (Retsch, Haan, Germany) and prepared as 25% (w/v) suspensions in DMEM supplemented with 10% newborn calf serum. Tissue homogenates were clarified by centrifugation at 14,000g for 10 min at 4°C, and the resulting supernatants were used for virus titration by plaque assay, as described previously (*23*). Details of experimental groups and sample sizes for mouse virus challenge experiments are provided in the corresponding figure legends. Mice were randomly assigned to cages, after which cages were randomly allocated to experimental groups.

### X-ray crystallography

For the ch.005 Fab structure, Fab in 1x TBS was concentrated down to ∼12 and 16 mg mL^−1^ using an Amicon spin concentrator with a 10 kDa MWCO (Millipore, cat. no. UFC9010). For the ls.019-peptide structure, the lower stalk (LS) peptide was slowly diluted into TBS for a final concentration of 1% DMSO; ls.019 scFv in 1x TBS was combined with the diluted peptide at a 1.5 molar excess of peptide to scFv and incubated overnight at room temperature (20-25°C). The scFv-peptide complex was concentrated to ∼12.2 mg mL^−1^ using an Amicon spin concentrator with a 3 kDa MWCO (Millipore, cat. no. UFC500324). Crystallization screens were set up using the sitting drop vapor diffusion method with equal volumes of concentrated protein and reservoir solution (100 nL each); drops were set using a Mosquito robot (SPT Labtech) and commercially available 96-well crystallization screens (Hampton Research, Rigaku, and Morpheus). Crystals were grown at room temperature (20-25°C) and observed in multiple conditions. For the ch.005 Fab structure, the condition that yielded the crystal harvested for structure determination consisted of 20% w/v polyethylene glycol monomethyl ether (PEG MME) 5,000, 0.2 M sodium formate, and 0.1 M bicine pH 8.5. For the ls.019-LS structure, the condition was composed of 0.2 M ammonium sulfate, 0.1 M 2-Morpholinoethanesulfonic acid (MES) monohydrate pH 6.5, and 30% w/v PEG MME 5,000. Upon looping, no cryoprotectants were added before cryocooling in liquid nitrogen.

X-ray diffraction data for both structures were collected at the Stanford Synchrotron Radiation Lightsource (SSRL) beamline 12-2 with an Eiger2 XE 16M pixel array detector (Dectris) at a wavelength of 0.979 Å and a temperature of 100 K. Data processed for each dataset were obtained using a single crystal and were indexed and integrated using an automated processing pipeline of XDS(*92*), and then merged using Aimless(*93*) in CCP4(*94*). Molecular replacement with Phaser(*95*) in Phenix(*96*) was used for structure determination. Search models were chosen based on which had the closest match to the protein sequence using IMGT/3Dstructure-DB(*97*) and that had good starting statistics.

For the ch.005 Fab structure, PDBs 5I1C and 6PZH were used as search models for the V_H_ and C_H_ and V_L_ and C_L_, respectively; CDR3 loops were trimmed prior to molecular replacement. For the ls.019-peptide structure, the following chains were used as search models: V_H_ from PDB 5N2K with trimmed CDR_H3_, V_L_ from PDB 6OC7 with trimmed CDR_L3_. Peptide was not included in molecular replacement search models. Molecular coordinates were subsequently refined by employing iterative rounds of automated and manual refinement in Phenix(*96*) and Coot(*98*), respectively. See Table S3 for further detail on collection and refinement statistics.

### Cryo-EM sample preparation

For SARS-CoV-2 SS – Fab complexes, purified SARS-CoV-2 SS (6 mg mL^−1^) was incubated with 1.1 molar excess of Fab for 30 min in 1x TBS. Quantifoil Cu R1.2/1.3 300 mesh grids (Electron Microscopy Sciences) were glow-discharged for 60 s at 10 mA on an easiGlow (PELCO) before sample application. Immediately prior to depositing 3.1 uL of the complex onto the glow-discharged grids, fluorinated octyl-maltoside (Anatrace) was added to the sample to a final concentration of 0.0084% w/v. Using a Vitrobot Mark IV (Thermo Fisher Scientific), grids were blotted for 3.5 s at room temperature, 100% humidity, and with blot force 3 or 4. They were then plunge-frozen in liquid ethane and transferred to liquid nitrogen for storage.

### Cryo-EM data collection and processing

Single-particle cryo-EM datasets were collected on a Titan Krios transmission electron microscope (Thermo Fisher Scientific) operated at 300 kV and equipped with either a K3 direct electron director (Gatan) for SARS-CoV-2-SS-ch.005 complex or Falcon4i direct electron director (Thermo Fisher Scientific) for all other complexes. Movie collection was automated using Smart EPU (Thermo Fisher Scientific) and followed an aberration-free image shift acquisition (AFIS) pattern with three shots per grid hole, and a total dose of 50 electrons/Å^2^ accumulated across 50 frames with calibrated pixel sizes of 0.85Å (SARS-CoV-2-SS-ch.005) or 0.92Å.

Following data collection, movies from all datasets underwent patch motion correction, patch contrast transfer function (CTF) estimation, reference-free particle picking, and particle extraction using cryoSPARC v4.6.2(*99*). A subset of particles was extracted, down sampled by 4x, and then iteratively refined using two-dimensional (2D) classification in cryoSPARC. 2D classes were then selected and used for template-based picking on the entire dataset. These particles, down sampled by 4x, were used to generate *ab initio* reconstructions in cryoSPARC, followed by heterogeneous refinement of the entire particle stack to generate an initial reference model. Particles underwent iterative rounds of 2D and three-dimensional (3D) classifications in cryoSPARC and Relion(*100*), followed by re-extraction of final particle stack without binning to be used as the input for non-uniform refinement in cryoSPARC.

For local refinement of the Fab-SS protomer interface, particles for the C3 non-uniform refinement jobs were symmetry expanded in cryoSPARC and used as input for the local refinement job in cryoSPARC. A mask of the V_H_ and V_L_ domains and the top of the SS protomer was made and used, along with applying C1 as symmetry. The mask used for the ch.005-SS local refinement job included the entire Fab domain, along with the top half of the SS protomer. The resolutions of all cryo-EM maps were estimated using the 0.143 Fourier shell correlation (FSC) cutoff in cryoSPARC. See Table S5 for further detail on collection and refinement statistics.

### Modeling and refinement of cryo-EM structures

Sharpened maps corresponding to the global C3 non-uniform refinement or the C1 local refinement were used for model building and refinement. Unsharpened maps were only used for building in N-linked glycans. Initial model coordinates for each protein were obtained by docking individual chains from homology models, identified using IMGT/3Dstructure-DB (*97*), (Table S4) into the sharpened cryo-EM maps using UCSF ChimeraX v1.9 (*101*). Sequences of the docked models were then updated manually in Coot (*98*) to align with the expressed protein. Models were then refined by employing iterative rounds of refinement in Phenix (*96*) and manual building and refinement in Coot. Refined model coordinates were ultimately validated in using MolProbity (*102*).

UCSF ChimeraX v1.9 (*101*) were used to visualize structures and create figures. Buried surface area (BSA), residue assignment to the epitopes and paratopes, and potential hydrogen bond assignments described were calculated using a 1.4 Å probe in PDBePISA v1.52 (*78, 79*). Antibody residue numbering was done in accordance with the Kabat (*103*) designation; CDRs were defined in accordance with IMGT (*104, 105*) designation.

### Quantification and statistical analyses

In flow cytometry binding experiments (Figure S2), a two-tailed paired t-test analysis (* p < 0.05, ** p < 0.01, *** p < 0.001, and **** p <0.0001) was performed on the geometric mean fluorescent intensity (gMFI, n=3 experiments). Statistical animal experiment analyses were performed using GraphPad Prism (GraphPad Software, Boston, MA, USA). For comparisons of body weight among experimental groups (Figure 5B), statistical significance was assessed at 5 days post-infection using one-way analysis of variance (ANOVA) followed by Tukey’s multiple-comparisons test. Survival distributions (Figure 5C) were compared using the log-rank (Mantel–Cox) test. Lung viral titers (Figure 5D) were compared using the Kruskal–Wallis test followed by Dunn’s multiple-comparisons test. Statistical significance is indicated as follows: * p < 0.05, ** p< 0.01, and *** p< 0.001; ns, not significant.

## ACKNOWLEDGEMENTS

We are indebted to all study participants and personnel of the Clinica Luganese Moncucco. We further thank Drs. T. Hatziioannou and P. Bieniasz (Rockefeller University) for sharing plasmids and protocols for work with SARS-CoV-2 pseudovirus. We thank members from the laboratory of Dr. P. Bjorkman (California Institute of Technology) for some of the SARS-CoV-2 spike protein expression constructs. We thank the Macromolecular Structure Group at the Nucleus, Stanford ChEM-H institute, especially Dr. D. Fernandez for support with protein crystal harvest and data collection. X-ray crystallography data for this work was collected at the Stanford Synchrotron Radiation Lightsource, SLAC National Accelerator Laboratory, which is supported by the U.S. Department of Energy, Office of Science, Office of Basic Energy Sciences under Contract No. DE-AC02-76SF00515. We thank Drs. M. Abernathy and S. Vasquez and Z. Contejean in Dr. C.O. Barnes’ lab (Stanford) for their advice with X-ray crystallography modeling and refinement, as well as at IRB Tristan Gasparetto and Andrea Celoria (for technical assistance), and Drs. Mattia Pedotti, Elia Tamagnini and Luca Varani (for SARS-CoV-2 spike trimer and soluble ACE2 reagents). Cryo-EM data for this work was collected at the SLAC National Accelerator Laboratory with assistance from Drs. B. Singal, H. Wang, and D. Fernandez-Martinez of the Stanford University Cryo-electron Microscopy Center (cEMc). We thank M. Dvorakova and M. Zemanova from Palus’ lab for technical assistance. The following reagent was obtained through BEI Resources, NIAID, NIH: SARS-related coronavirus 2, mouse-adapted, MA10 variant (in isolate USA-WA1/2020 backbone), infectious clone (ic2019-nCoV MA10) in Vero TMPRSS2/ACE2, NR-60440, contributed by Ralph S. Baric. Finally, we thank all members of the Robbiani and Barnes labs for their careful review and feedback on the manuscript.

## AUTHOR CONTRIBUTIONS

**Conceptualization:** AAR, VC, CG, MP, DFR, COB

**Methodology:** AAR, VC, TCR, CG, MP, DFR, COB

**Investigation:** AAR, VC, MP, TCR, MEA, MKP, DV, YEL, ME, JC, SM, DJ, CG, BC, MM, VG, MB, VC, AFP, CG, AC

**Visualization:** AAR, VC, MP

**Funding acquisition:** MP, DFR, COB

**Project administration**: CG, MP, DFR, COB

**Supervision:** CG, MP, DFR, COB

**Writing – original draft:** AAR, COB

**Writing – review & editing**: AAR, MP, DFR, COB

## DATA AND MATERIALS AVAILABILITY

All expression plasmids generated in this study for CoV spike proteins, human Fabs and IgGs are available upon request and may require a Materials Transfer Agreement (MTA). The atomic models generated via X-ray crystallography for the ch.005 Fab and ls.019-LS peptide complex have been deposited to the Protein Data Bank (PDB) under accession codes: 10QO and 10NI, respectively. The atomic models and cryo-EM maps generated for ch.005-SS (global refinement), ch.005-SS (local refinement), ch.007-SS (global refinement), ch.007-SS (local refinement), and ch.010-SS (local refinement) have been deposited to the PDB and the Electron Microscopy Databank (EMDB) under accession codes PDB: 9ZLQ, 9ZLR, 9ZLS, 9ZLT, and 9ZLU and EMD: −74407, −74408, −74409, −74410, and −74412. The cryo-EM maps generated for ch.010-SS (global refinement) have been deposited to the EMDB under accession code EMD-74411. The script for coldspot identification is available at GitHub (https://github.com/cavallilab/coldspot). Any additional information required to reanalyze the data reported in this paper or requests for resources and reagents should be directed to and will be fulfilled by the lead contact, Dr. Christopher O. Barnes.

## Supplementary materials

**Table S1.**
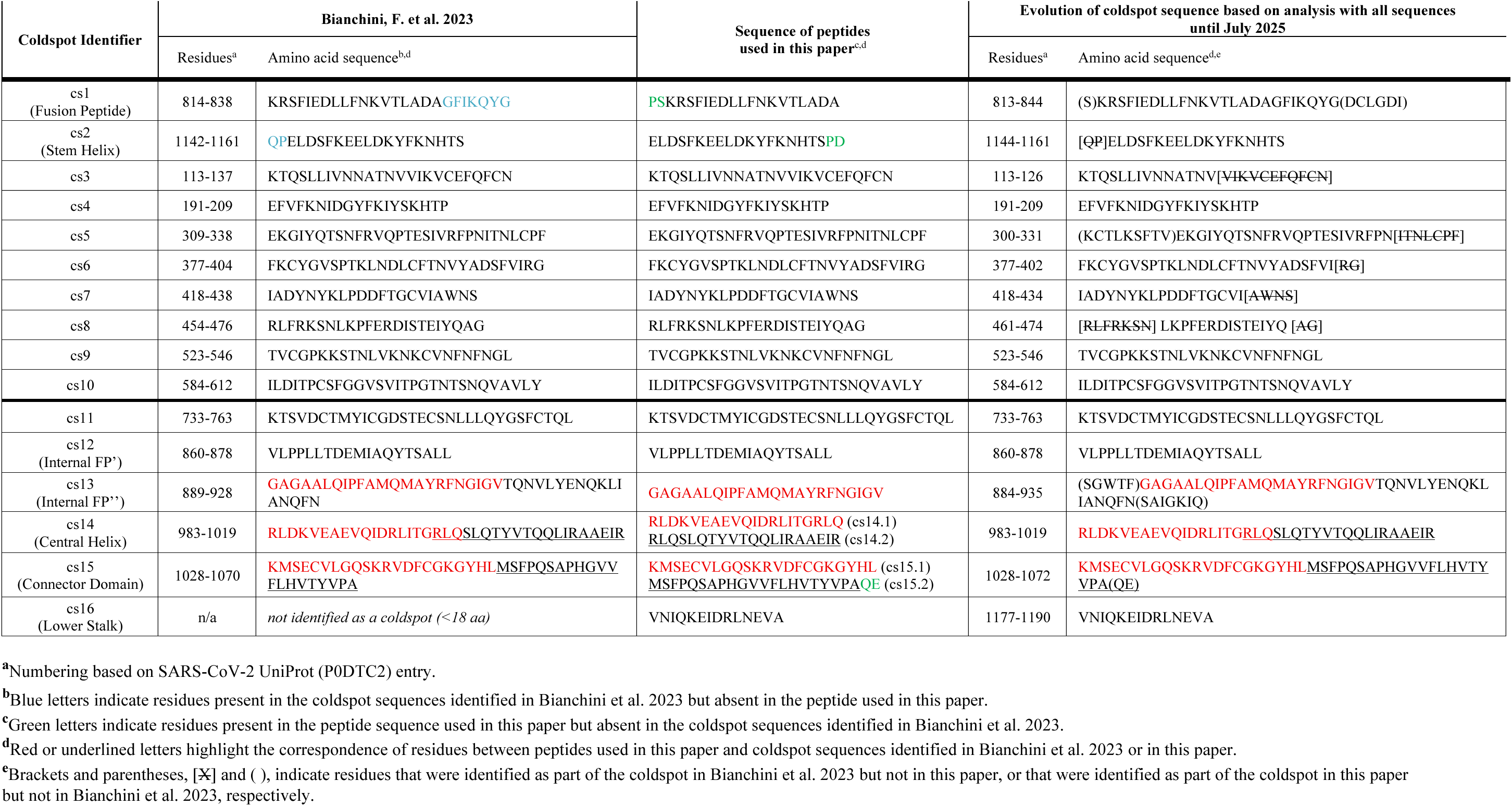
Amino acid sequence of coldspots and corresponding peptides. Related to Figure 1.

**Table S2.** Binding kinetics of CH and LS monoclonal antibodies. Related to Figure 2.

| Antibody | Antigen | $k_{on} (M^{-1}s^{-1})$ | $k_{off} (s^{-1})$ | $K_d (nM)$ |
| --- | --- | --- | --- | --- |
| **ch.005** | SARS-CoV-2 SS | $3.6 \times 10^5$ | $3.6 \times 10^{-4}$ | 1.0 |
| | SARS-CoV-2 D994A SS | $2.6 \times 10^2$ | $6.1 \times 10^{-2}$ | $2.3 \times 10^6$ |
| | SARS-CoV-2 R995A SS | $2.2 \times 10^5$ | $5 \times 10^{-4}$ | 2.3 |
|  | SARS-CoV-2 Q1002A SS | N.D. | N.D. | N.D. |
| **ch.007** | SARS-CoV-2 SS | $1.4 \times 10^5$ | $1.5 \times 10^{-4}$ | 1.1 |
| | SARS-CoV-2 D994A SS | $4.7 \times 10^3$ | $1.5 \times 10^{-3}$ | 317 |
|  | SARS-CoV-2 R995A SS | N.D. | N.D. | N.D. |
| | SARS-CoV-2 Q1002A SS | $3.6 \times 10^4$ | $5.7 \times 10^{-4}$ | 16 |
| **ch.010** | SARS-CoV-2 SS | $3.7 \times 10^5$ | $5.3 \times 10^{-4}$ | 1.4 |
| | SARS-CoV-2 D994A SS | $2.1 \times 10^4$ | $1.7 \times 10^{-2}$ | 802 |
| | SARS-CoV-2 R995A SS | $5.3 \times 10^5$ | $4.5 \times 10^{-3}$ | 8.4 |
| | SARS-CoV-2 Q1002A SS | $1.5 \times 10^5$ | $5.4 \times 10^{-4}$ | 3.4 |
| | MERS-CoV SSv2 | $1.5 \times 10^5$ | $1.7 \times 10^{-2}$ | 115 |
| **54043-5<sup>a</sup>** | MERS-CoV SSv2 | $2.1 \times 10^5$ | $2.2 \times 10^{-3}$ | 10 |
| **ls.015** | SARS-CoV-2 spike 6P | $2.2 \times 10^4$ | $3.2 \times 10^{-3}$ | 148 |
| **ls.019** | SARS-CoV-2 spike 6P | $1.1 \times 10^4$ | $1.3 \times 10^{-3}$ | 114 |

<sup>a</sup>Johnson, N.V. *et al.* 2024<sup>b</sup>N.D., not determined

**Table S3.** X-ray crystallography data collection and refinement statistics. Related to Figure 2 and 3.

|  | ch.005 Fab | Is.019 scFv-LS peptide |
| --- | --- | --- |
| PDB ID | 1OQO | 1ONI |
| **Data collection<sup>a</sup>** |  |  |
| Space group | P 41 | P 21 21 2 |
| Unit cell (Å) | 69.545, 69.545, 86.108 | 133.257, 38.765, 47.774 |
| $\alpha$, $\beta$, $\gamma$ (°) | 90, 90, 90 | 90, 90, 90 |
| Wavelength (Å) | 0.979 | 0.979 |
| Resolution (Å) | 36.61 – 1.6 (1.63 – 1.6) | 38.83 – 1.788 (1.81 – 1.79) |
| Unique Reflections | 53,716 | 23,663 |
| Completeness (%) | 99.4 (92.9) | 99.2 (94.0) |
| Redundancy | 7.3 (4.0) | 4.4 (4.2) |
| CC<sub>1/2</sub> (%) | 0.998 (0.794) | 0.992 (0.352) |
| I / σI | 14.8 (2.6) | 5.2 (0.6) |
| Mosaicity (°) |  | 0.24 |
| R<sub>merge</sub> (%) | 0.066 (0.612) | 0.109 (1.56) |
| R<sub>pim</sub> (%) | 0.038 (0.467) | 0.080 (1.14) |
| Wilson B-factor | 18.76 | 26.37 |
| **Refinement and Validation** |  |  |
| Resolution (Å) | 1.6 | 1.79 |
| Number of atoms |  |  |
| Protein | 6,031 | 1,741 |
| Ligand | 0 | 0 |
| Waters | 34 | 57 |
| R<sub>work</sub>/R<sub>free</sub> (%) | 0.2112 / 0.2279 | 0.2159 / 0.2473 |
| R.m.s. deviations |  |  |
| Bond lengths (Å) | 0.005 | 0.002 |
| Bond angles (°) | 0.799 | 0.61 |
| MolProbity score | 0.89 | 1.62 |
| Clashscore (all atom) | 0.5 | 2.36 |
| Poor rotamers (%) | 1.17 | 2.75 |
| Ramachandran plot |  |  |
| Favored (%) | 97.09 | 96.05 |
| Allowed (%) | 2.91 | 3.51 |
| Disallowed (%) | 0.00 | 0.44 |
| Average B-factor (Å) | 26.67 | 36.3 |

<sup>a</sup>Numbers in parentheses correspond to the highest resolution shell

**Table S4.**
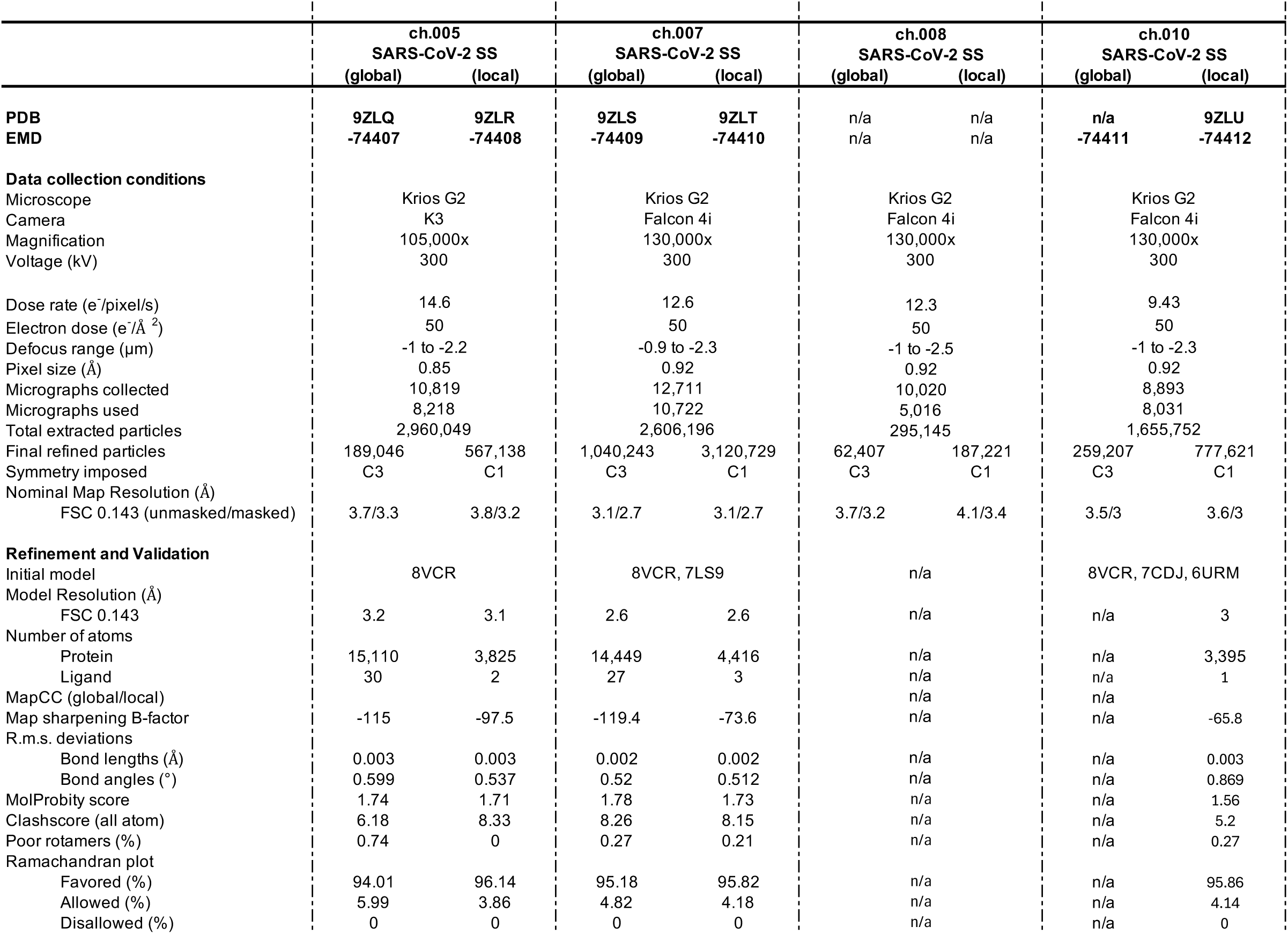
Cryo-EM data collection and refinement statistics. Related to Figure 3 and 4.

|  | ch.005<br>SARS-CoV-2 SS |  | ch.007<br>SARS-CoV-2 SS |  | ch.008<br>SARS-CoV-2 SS |  | ch.010<br>SARS-CoV-2 SS |  |
| --- | --- | --- | --- | --- | --- | --- | --- | --- |
|  | (global) | (local) | (global) | (local) | (global) | (local) | (global) | (local) |
| **PDB** | **9ZLQ** | **9ZLR** | **9ZLS** | **9ZLT** | n/a | n/a | **n/a** | **9ZLU** |
| **EMD** | **-74407** | **-74408** | **-74409** | **-74410** | n/a | n/a | **-74411** | **-74412** |
| **Data collection conditions** |  |  |  |  |  |  |  |  |
| Microscope | Krios G2 |  | Krios G2 |  | Krios G2 |  | Krios G2 |  |
| Camera | K3 |  | Falcon 4i |  | Falcon 4i |  | Falcon 4i |  |
| Magnification | 105,000x |  | 130,000x |  | 130,000x |  | 130,000x |  |
| Voltage (kV) | 300 |  | 300 |  | 300 |  | 300 |  |
| Dose rate (e<sup>-</sup>/pixel/s) | 14.6 |  | 12.6 |  | 12.3 |  | 9.43 |  |
| Electron dose (e<sup>-</sup>/Å<sup>2</sup>) | 50 |  | 50 |  | 50 |  | 50 |  |
| Defocus range (µm) | -1 to -2.2 |  | -0.9 to -2.3 |  | -1 to -2.5 |  | -1 to -2.3 |  |
| Pixel size (Å) | 0.85 |  | 0.92 |  | 0.92 |  | 0.92 |  |
| Micrographs collected | 10,819 |  | 12,711 |  | 10,020 |  | 8,893 |  |
| Micrographs used | 8,218 |  | 10,722 |  | 5,016 |  | 8,031 |  |
| Total extracted particles | 2,960,049 |  | 2,606,196 |  | 295,145 |  | 1,655,752 |  |
| Final refined particles | 189,046 | 567,138 | 1,040,243 | 3,120,729 | 62,407 | 187,221 | 259,207 | 777,621 |
| Symmetry imposed | C3 | C1 | C3 | C1 | C3 | C1 | C3 | C1 |
| Nominal Map Resolution (Å) |  |  |  |  |  |  |  |  |
| FSC 0.143 (unmasked/masked) | 3.7/3.3 | 3.8/3.2 | 3.1/2.7 | 3.1/2.7 | 3.7/3.2 | 4.1/3.4 | 3.5/3 | 3.6/3 |
| **Refinement and Validation** |  |  |  |  |  |  |  |  |
| Initial model | 8VCR |  | 8VCR, 7LS9 |  | n/a |  | 8VCR, 7CDJ, 6URM |  |
| Model Resolution (Å) |  |  |  |  |  |  |  |  |
| FSC 0.143 | 3.2 | 3.1 | 2.6 | 2.6 | n/a |  | n/a | 3 |
| Number of atoms |  |  |  |  |  |  |  |  |
| Protein | 15,110 | 3,825 | 14,449 | 4,416 | n/a |  | n/a | 3,395 |
| Ligand | 30 | 2 | 27 | 3 | n/a |  | n/a | 1 |
| MapCC (global/local) |  |  |  |  | n/a |  | n/a |  |
| Map sharpening B-factor | -115 | -97.5 | -119.4 | -73.6 | n/a |  | n/a | -65.8 |
| R.m.s. deviations |  |  |  |  |  |  |  |  |
| Bond lengths (Å) | 0.003 | 0.003 | 0.002 | 0.002 | n/a |  | n/a | 0.003 |
| Bond angles (°) | 0.599 | 0.537 | 0.52 | 0.512 | n/a |  | n/a | 0.869 |
| MolProbity score | 1.74 | 1.71 | 1.78 | 1.73 | n/a |  | n/a | 1.56 |
| Clashscore (all atom) | 6.18 | 8.33 | 8.26 | 8.15 | n/a |  | n/a | 5.2 |
| Poor rotamers (%) | 0.74 | 0 | 0.27 | 0.21 | n/a |  | n/a | 0.27 |
| Ramachandran plot |  |  |  |  |  |  |  |  |
| Favored (%) | 94.01 | 96.14 | 95.18 | 95.82 | n/a |  | n/a | 95.86 |
| Allowed (%) | 5.99 | 3.86 | 4.82 | 4.18 | n/a |  | n/a | 4.14 |
| Disallowed (%) | 0 | 0 | 0 | 0 | n/a |  | n/a | 0 |

**Figure S1.**
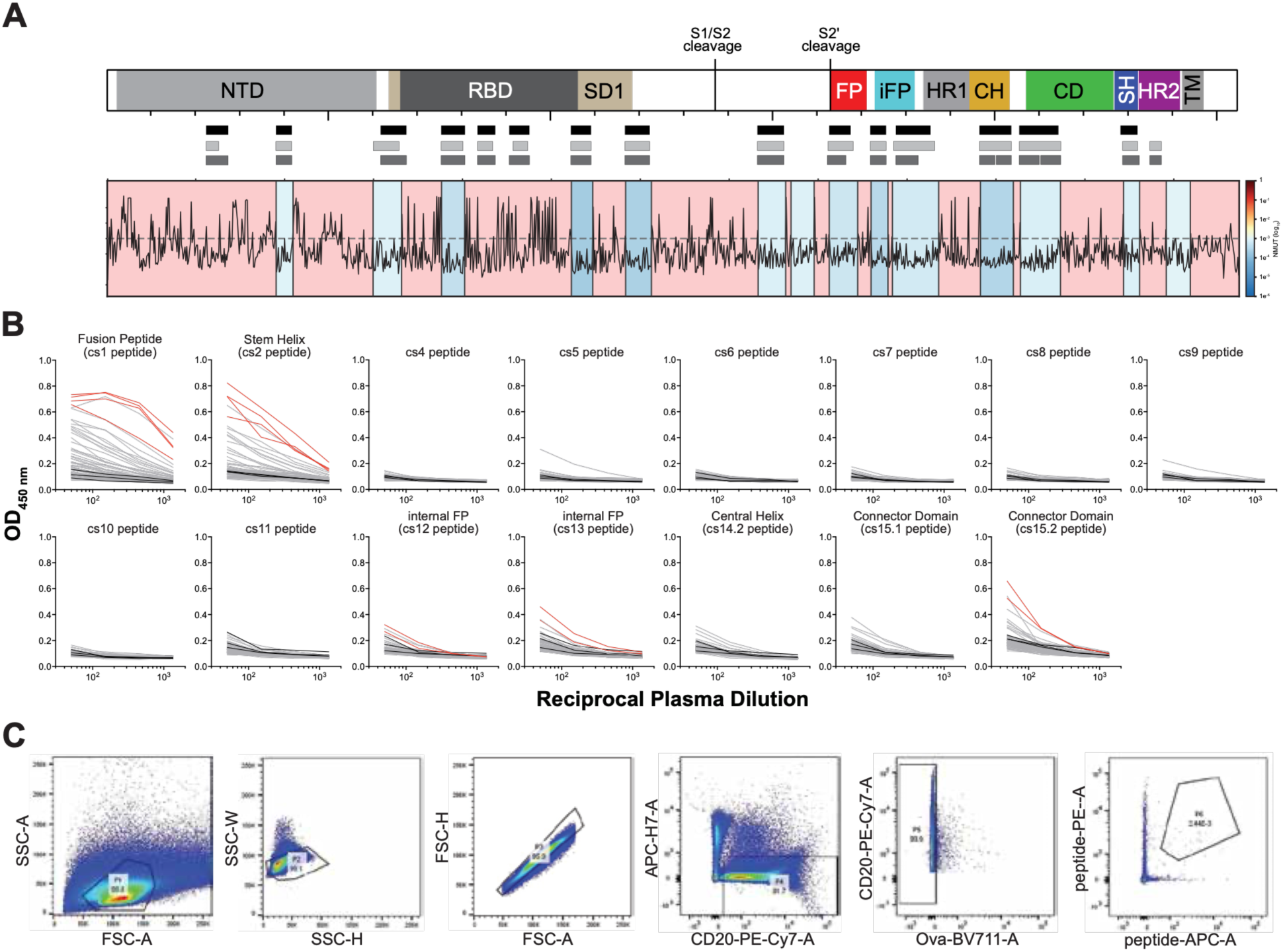
Identification of coldspot antibodies to the SARS-CoV-2 spike internal fusion peptide, central helix, connector domain, and lower stalk, related to Figure 1. **(A)** Top: Cartoon diagram of the functional domains of the SARS-CoV-2 spike protein. Coldspot regions and corresponding peptides are shown with rectangles below the diagram as follows: top row, coldspots as identified in Bianchini et al (black); middle row, evolution of the coldspots from Bianchini et al. by taking into consideration of sequences until July 2025 (light grey); bottom row, coldspot peptides used in this paper (dark grey). See also Table S1. Bottom: mapping of the frequency of amino acid changes (until July 2025). **(B)** ELISA plots for plasma IgG reactivity to coldspot peptides in spike. Each line represents a convalescent individual; red lines indicate donors selected for antibody isolation; black lines indicate prepandemic controls. Note: a peptide corresponding to cs3 could not be synthesized. **(C)** Representative gating strategy for sorting S2 coldspot peptide-specific B cells by flow cytometry.

**Figure S2.**
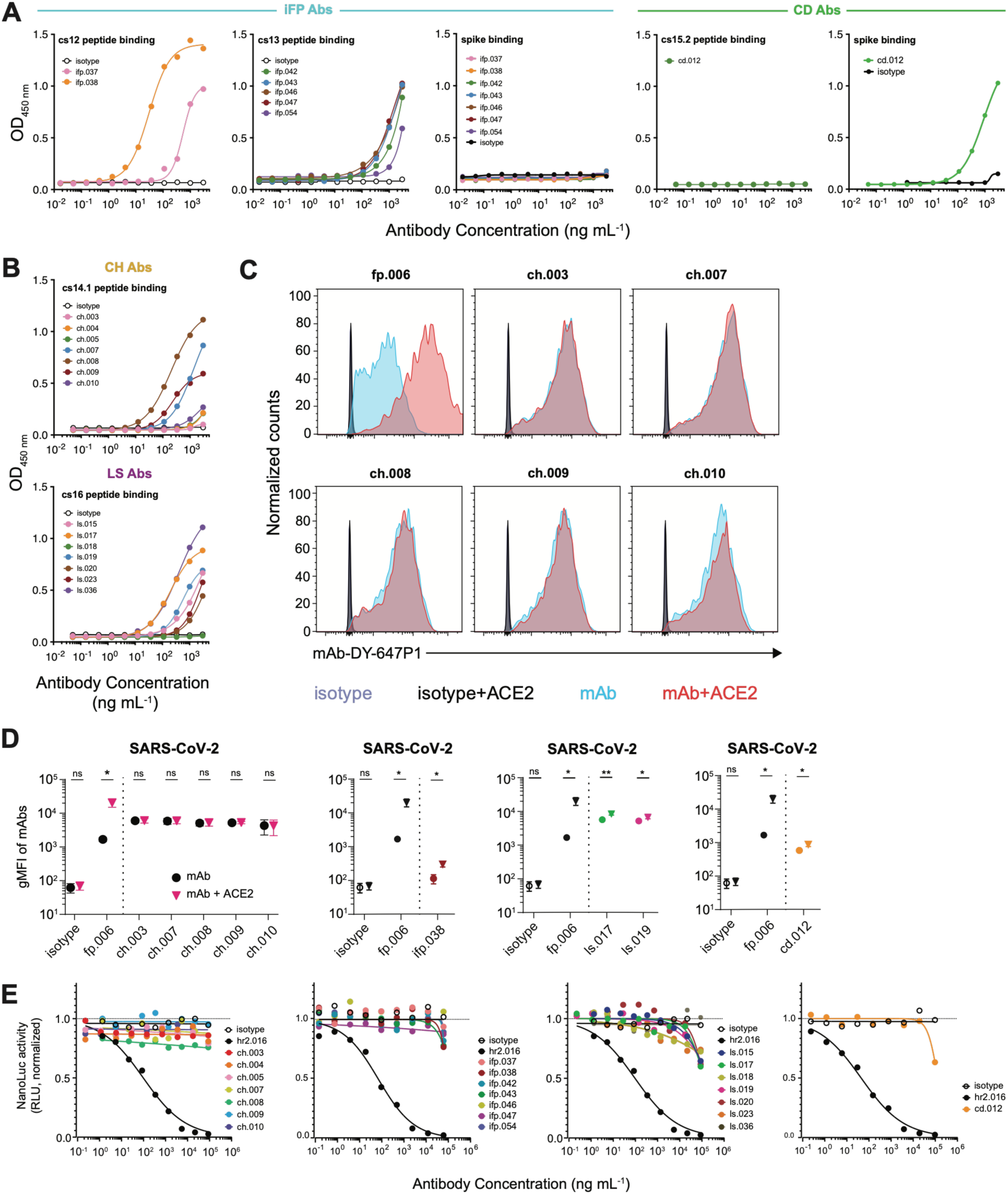
Profiling of coldspot monoclonal antibodies’ binding and neutralization of SARS-CoV-2 pseudovirus, related to Figure 1. **(A-B)** ELISAs measuring recombinant monoclonal antibody binding to (A) cognate peptide and SARS-CoV-2 spike for iFP (cs12 and cs13) and CD (cs15.2) and (B) CH (cs14.1) and LS (cs16) peptides. See Figure 1E for comparative binding of CH and LS antibodies to full-length spike. **(C)** Representative flow cytometry plots of anti-CH antibodies binding to SARS-CoV-2 spike expressed on HEK293 cells, in the presence or absence of soluble human ACE2. Isotype and anti-FP antibody fp.006 are shown as references. **(D)** Quantification of the geometric mean fluorescence intensity (gMFI) for CH, iFP, CD, and LS antibodies against spike expressed on HEK293 cells, in the presence or absence of soluble human ACE2. **(E)** Represenative dose-response curves of antibody neutralizing capacity depicted as changes in normalized relative luminescence (RLU) in cell lysates 48h after infection with ancestral SARS-CoV-2 pseudovirus in the presence of increasing concentrations of CH, iFP, LS, and CD antibodies, from left to right, respectively.

**Figure S3.**
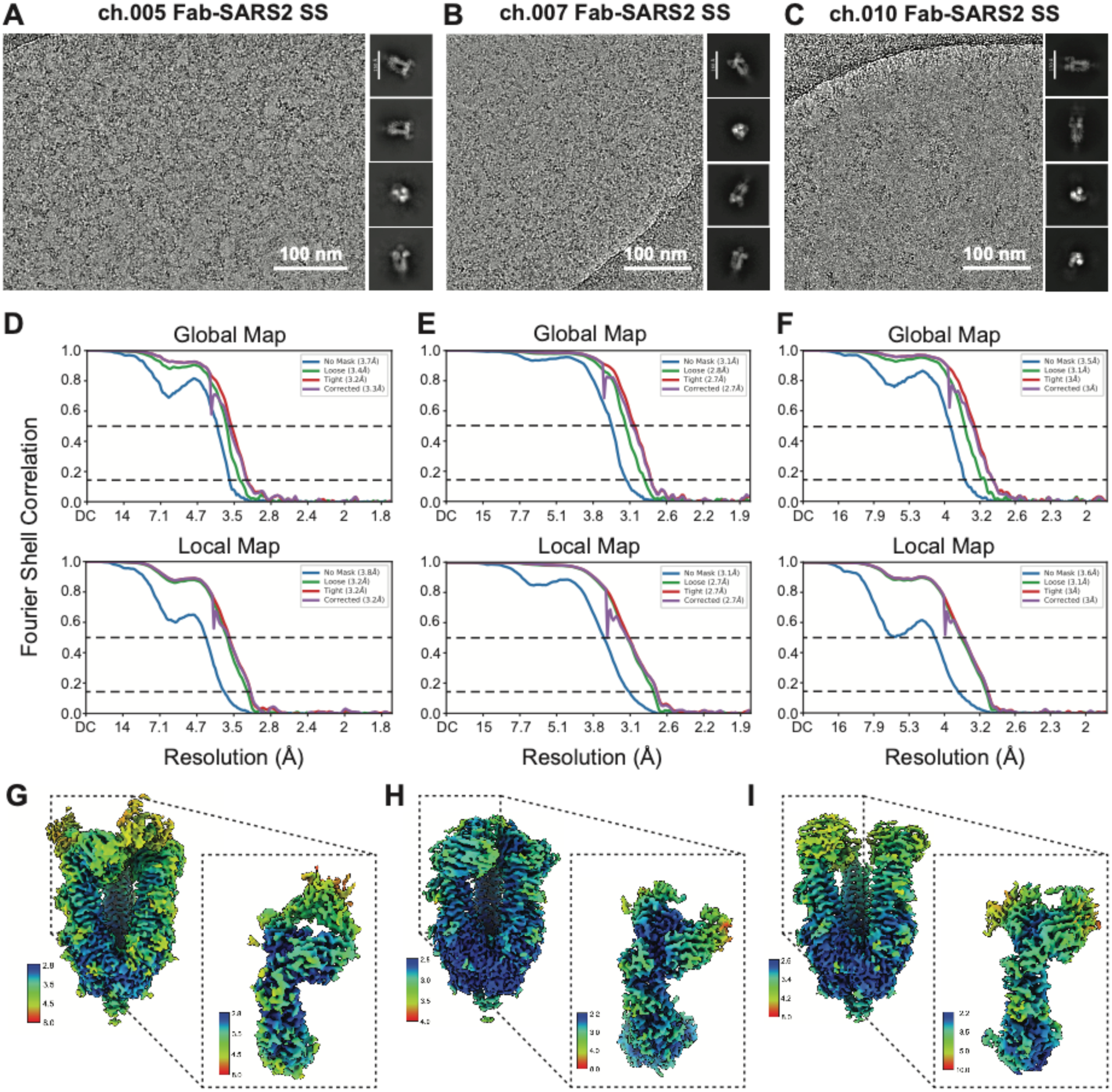
Cryo-EM data processing and validation of ch.005, ch.007, and ch.010 Fabs bound to SARS-CoV-2 stabilized stem (SS), related to Figure 3. **(A-C)** Representative micrographs and 2D class averages for **(A)** ch.005, **(B)** ch.007, and **(C)** ch.010 Fabs complexed with SARS-CoV-2 SS (see Table S4). **(D-F)** Gold-standard FSC plots for the global (top) or local (bottom) refinements for **(D)** ch.005-SS, **(E)** ch.007-SS, and **(F)** ch.010-SS complexes. **(G-I)** Local resolution estimations for **(G)** ch.005-SS, **(H)** ch.007-SS, and **(I)** ch.010-SS. Insets depict local resolution estimations for the Fab-protomer focused refinements.

**Figure S4.**
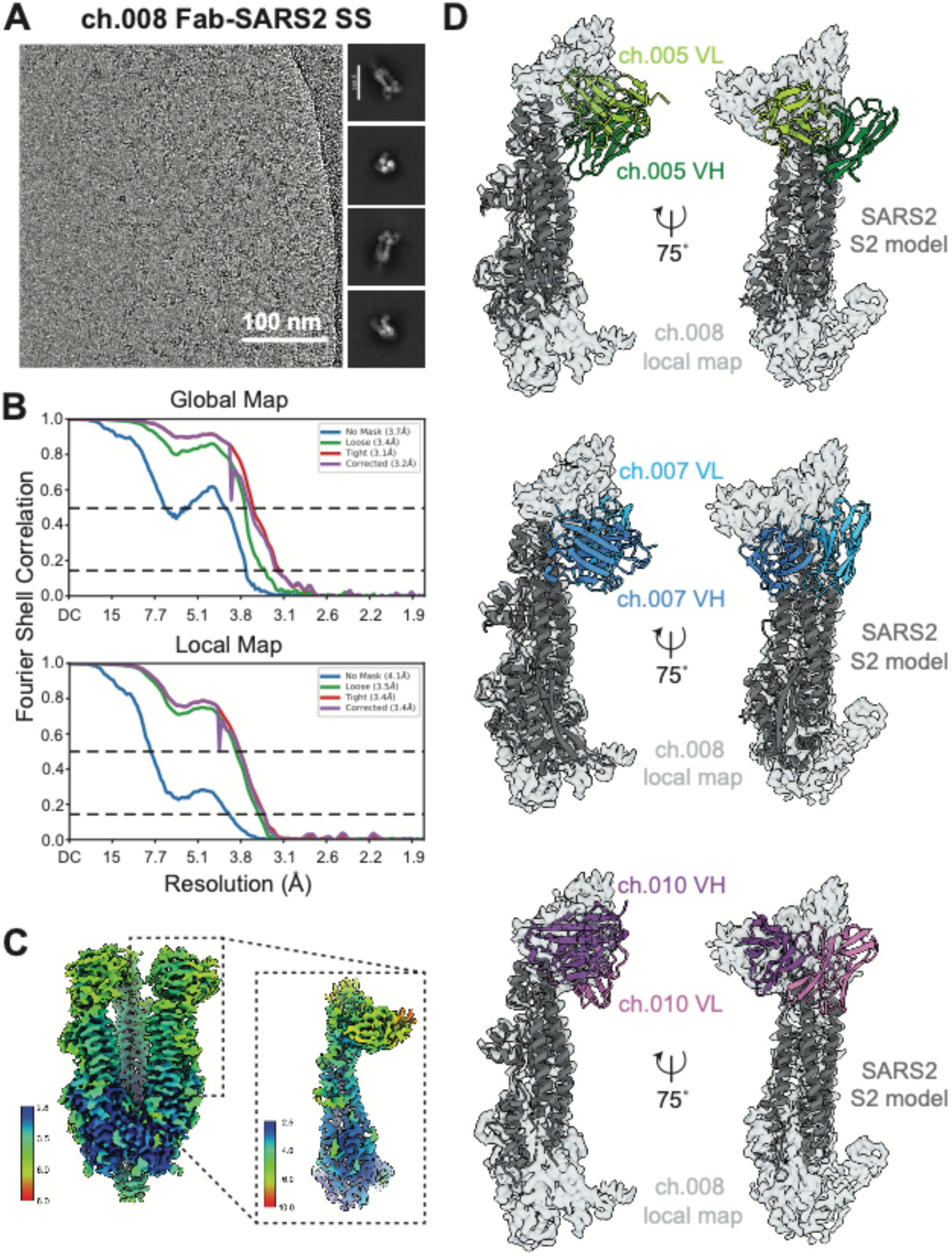
Cryo-EM data processing, validation, and characterization of ch.008, related to Figure 3. **(A)** Representative micrographs and 2D class averages for ch.008 complexed with SARS-CoV-2 SS (see Table S4). **(B)** Gold-standard FSC plots for the global (top) or local (bottom) refinements for ch.008-SS. **(C)** Local resolution estimations for ch.008-SS. Insets depict local resolution estimations for the Fab-protomer focused refinements. **(D)** Dockings of local refinement models of ch.005-SS (top), ch.005-SS (middle), and ch.010-SS (bottom) into the ch.008-SS local refinement map for comparison of binding pose.

**Figure S5.**
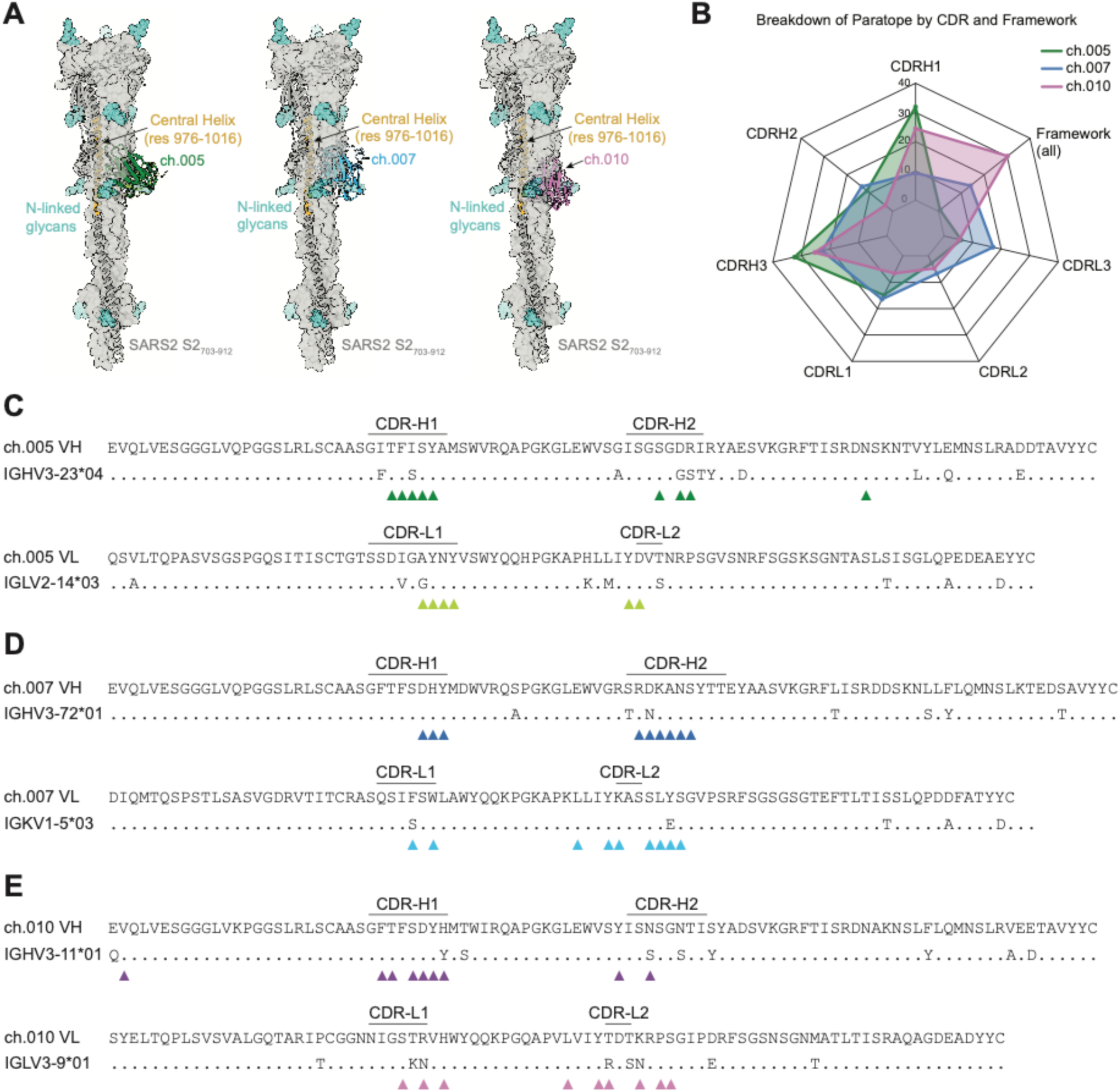
Characterization of epitope and paratope contacts for ch.005, ch.007, and ch.010, related to Figures 3 and 4. **(A)** Modeling of ch.005 (left), ch.007 (middle), and ch.010 (right) V_H_-V_L_ engaging their epitopes in the SARS-CoV-2 postfusion state (PDB 7E9T). Models were generated by aligning C*α* of residues 988-1028 of the local refinement models to the postfusion model. **(B)** Radar plot depicting the percent contribution of individual V_H_ and V_L_ CDR loops and total framework residues to the antibody paratopes for ch.005 (green), ch.007 (blue), and ch.010 (purple). **(C-E)** Sequence alignment of antibody V_H_ (top) or V_L_ (bottom) to their respective Ig germlines for ch.005 **(C)**, ch.007 **(D)**, and ch.010 **(E).** Dots represent agreement with germline. Residues involved with the paratope are denoted by triangles.

**Figure S6.**
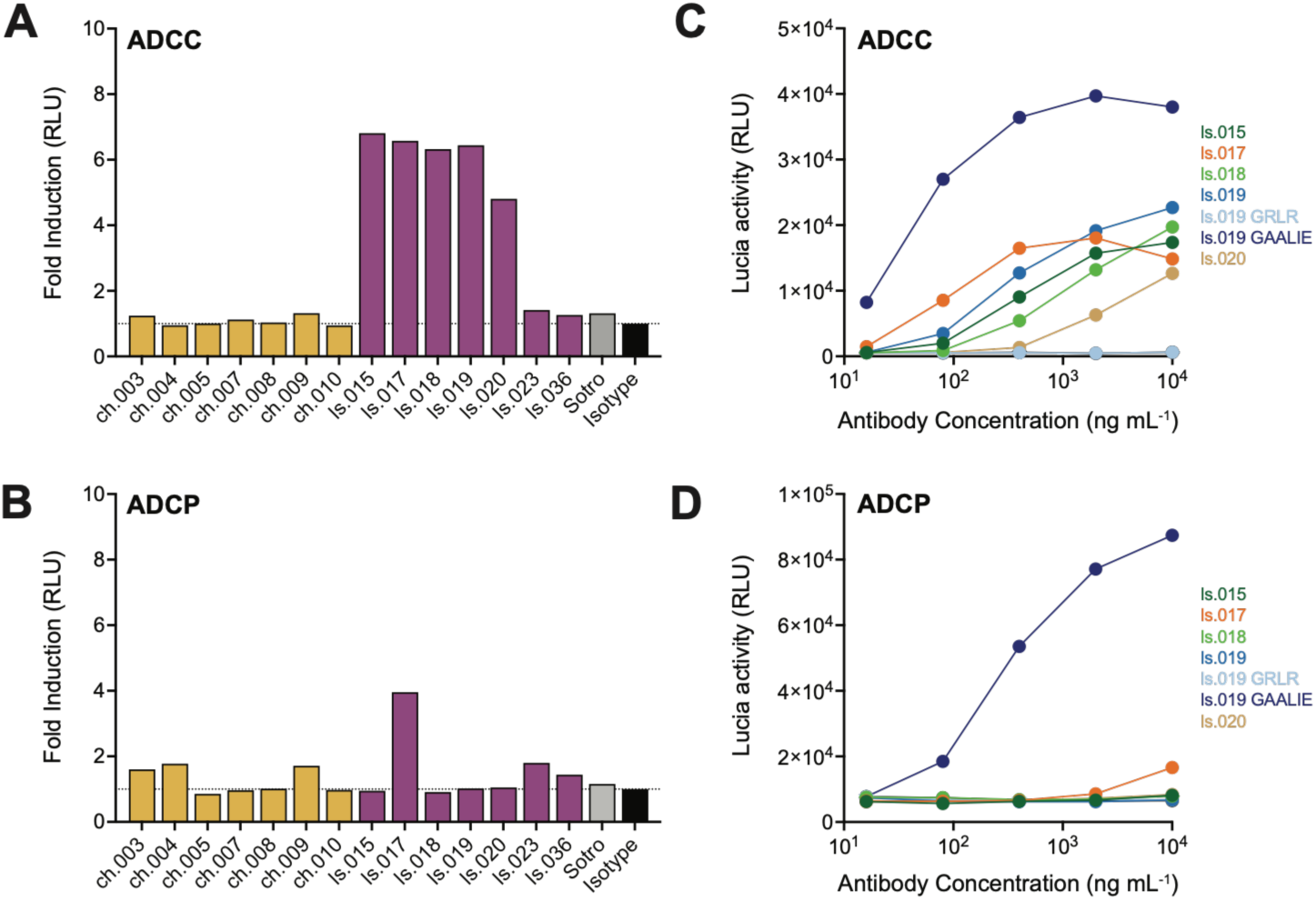
*In vitro* Fc-mediated effector functions by CH and LS antibodies, related to Figure 5. **(A-B)** FcγR reporter activation screen of CH and LS antibodies: FcγRIIIA/CD16A (ADCC) **(A)** and FcγRIIA/CD32A (ADCP) **(B)** reporter activation induced by antibodies in the presence of cells expressing SARS-CoV-2 spike. Lucia luciferase activity is shown as fold induction relative to the isotype control. Each bar represents an antibody tested at 10 μg/mL from a single experiment. Sotrovimab, Sotro, shown for reference. **(C-D)** Dose-dependent FcγR reporter activation by LS antibodies: FcγRIIIA/CD16A (ADCC) **(C)** and FcγRIIA/CD32A (ADCP) **(D)** reporter activation shown as raw relative light units (RLU) across five-fold serial antibody dilutions starting at 10 μg/mL. Each curve represents a monoclonal antibody. Data are representative of two independent experiments performed in duplicate.

**Figure S7.**
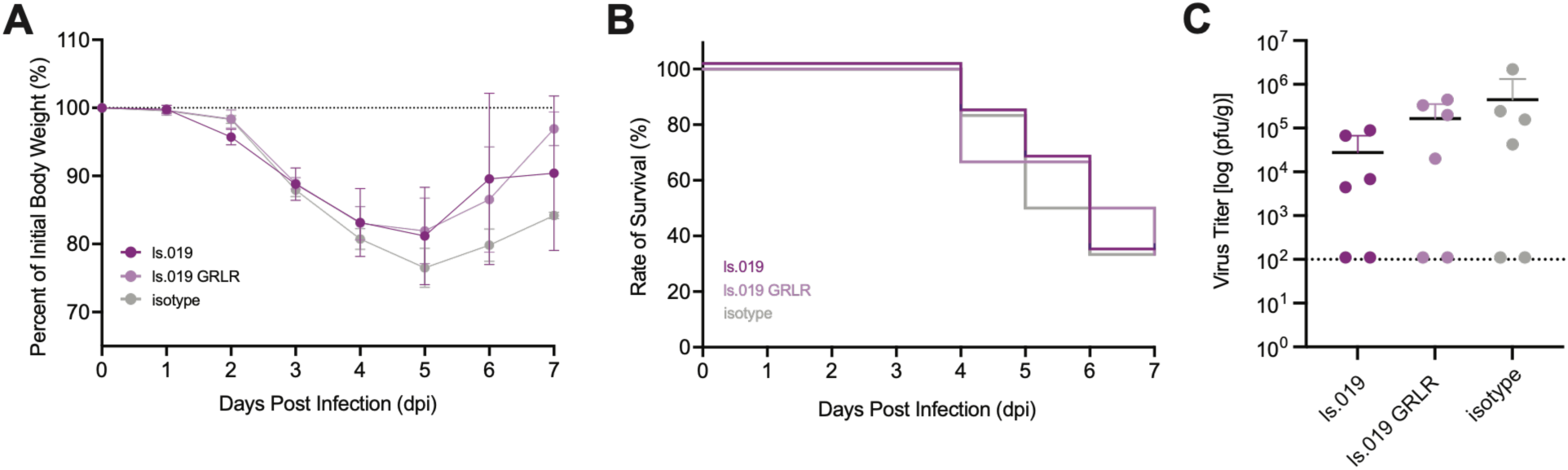
LS-directed antibody ls.019 does not confer protection *in vivo*, related to Figure 5. **(A)** Body weight curves for BALB/c mice treated prophylactically with 500 µg of ls.019 or ls.019 GRLR compared to mice treated with isotype control antibody (500 µg), prior to i.n. inoculation with SARS-CoV-2 MA10. n = 6 per group; error bars represent the standard error of the mean but statistical analysis was not performed. **(B-C)** Corresponding survival curve **(B)** and quantification of viral titer within lungs **(C)**, related to panel (A).

### Other supplementary material for the manuscript includes the following

**Data File S1.** Sequences of anti-coldspot antibodies.

**Data File S2.** Sequences and neutralizing and binding data of the monoclonal antibodies.

**Data File S3.** Alignment of the SARS-CoV-2 coldspot peptide sequences to related CoV species.

